# Discovery of lydiamycin A biosynthetic gene cluster in the plant pathogen *Rhodococcus fascians* guides structural revision and identification of molecular target

**DOI:** 10.1101/2024.11.13.623425

**Authors:** Jonathan J. Ford, Javier Santos-Aberturas, Edward S. Hems, Joseph W. Sallmen, Lena A. K. Bögeholz, Guy Polturak, Anne Osbourn, Joseph A. Wright, Marina V. Rodnina, Danny Vereecke, Isolde M. Francis, Andrew W. Truman

## Abstract

The natural products actinonin and matlystatin feature an *N*-hydroxy-2-pentyl-succinamyl (HPS) chemophore that facilitates metal chelation and confers their metalloproteinase inhibitory activity. Actinonin is the most potent natural inhibitor of peptide deformylase (PDF) and exerts antimicrobial and herbicidal bioactivity by disrupting protein synthesis. Here, we used a genomics-led approach to identify candidate biosynthetic gene clusters (BGCs) hypothesised to produce novel HPS-containing natural products. We show that one of these BGCs is on the pathogenicity megaplasmid of the plant pathogen *Rhodococcus fascians* and produces lydiamycin A, a macrocyclic pentapeptide. The presence of genes predicted to make a HPS-like chemophore informed the structural recharacterisation of lydiamycin via NMR and crystallography to show it features a rare 2-pentyl-succinyl chemophore. We demonstrate that lydiamycin A inhibits bacterial PDF in vitro and show that a cluster-situated PDF gene confers resistance to lydiamycin A, representing a novel self-immunity mechanism associated with the production of a PDF inhibitor. *In planta* competition assays showed that lydiamycin enhances the fitness of *R. fascians* during plant colonisation. This study highlights how a BGC can inform the structure, biochemical target and ecological function of a natural product.

## INTRODUCTION

Natural products (NPs) provide producing organisms with beneficial traits, such as intra-species communication, nutrient acquisition, or growth inhibition of competing organisms^1^. NPs are also critical in medicine and agriculture, where they have diverse uses, including as immunosuppressive, antiparasitic, antifungal and anticancer medicines^2^, as well as accounting for the majority of approved antibiotics^3^. This success can be attributed to the evolutionarily beneficial selection of potent chemotypes that strongly bind relevant biological targets. Actinonin and matlystatin are structurally related actinobacterial NPs that feature the same *N*-hydroxy-(*R*)-2-pentyl-succinamyl (HPS) chemophore^4,5^ (Fig. 1A). This chemophore chelates metal ions in the active sites of metalloproteins via a hydroxamate group^6^. This activity means that actinonin exhibits nanomolar activity towards multiple targets that have been validated in disease models in mice, including as an anti-tumour candidate^7^, acute renal failure^8^, anti-malarial^9^, anti-obesity^10^ and anti-bacterial, where the HPS moiety is a structural mimic of *N*-formylmethionine^11^, the substrate of the metalloproteinase peptide deformylase (PDF). PDF catalyses the deformylation of the initiator methionine residue on the growing nascent peptide, which is necessary for protein maturation, and is essential for the viability of most bacteria. Synthetic metalloprotease inhibitors inspired by actinonin have also been investigated in clinical trials as antibiotics, including LBM415 and GSK1322322^12^.

**Figure 1.**
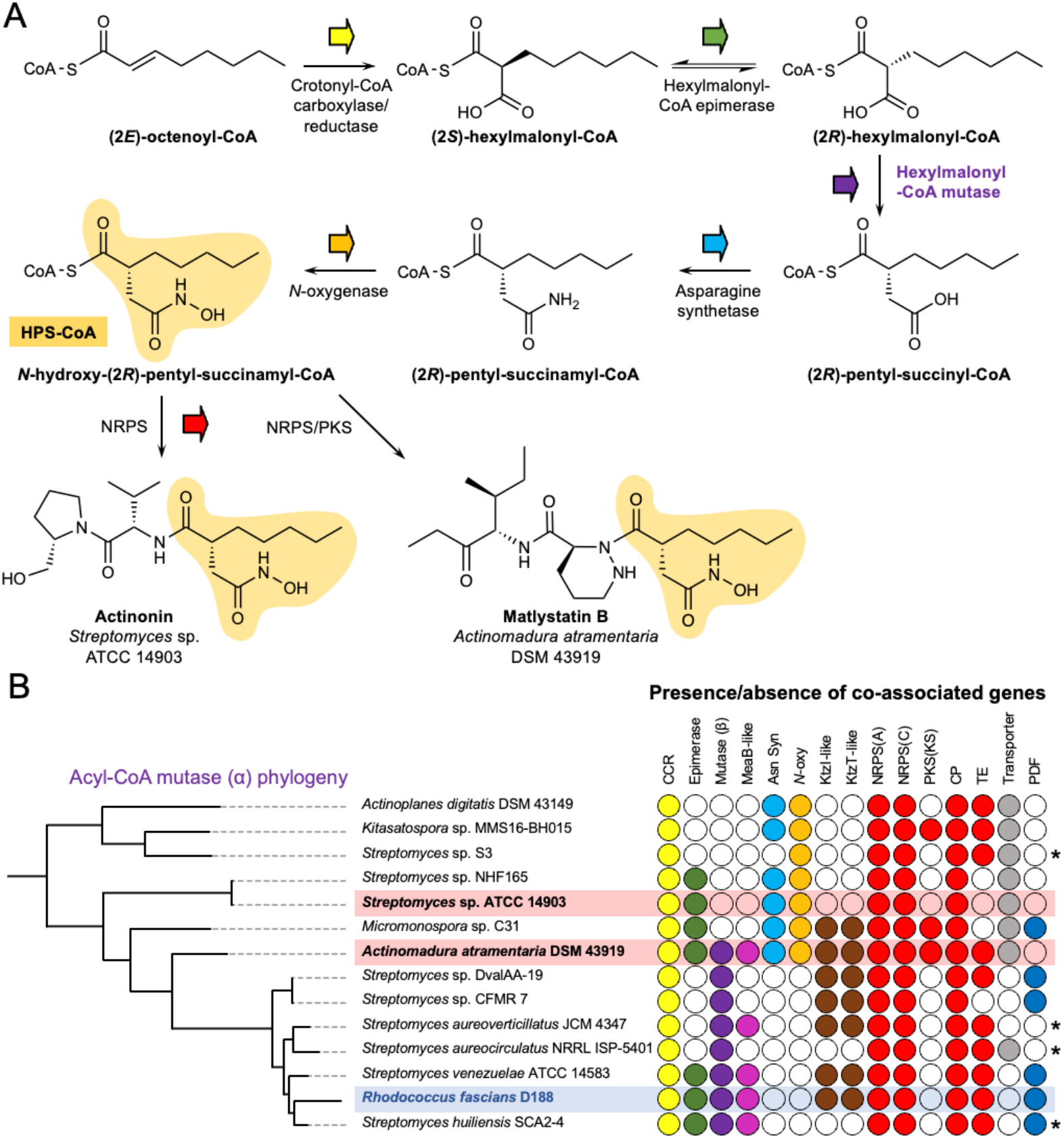
Mutase-guided BGC discovery. A. Proposed HPS biosynthesis for actinonin and matlystatin. B. Clade of actinobacterial mutases that are highly co-associated with the proposed HPS biosynthesis genes (colour-coded as in panel A), NRPS/PKS genes (red) and peptide deformylase gene (green). A filled circle indicates the presence of a homologous gene in proximity to the mutase gene. The actinonin and matlystatin producers are indicated in red. Asterisks indicate where BGCs are at the end of a contig and may therefore be incomplete. Abbreviations: CCR = crotonyl-CoA carboxylase-reductase, asparagine synthetase, *N*-oxygenase, NRPS(A) = NRPS adenylation domain, NRPS(C) = NRPS condensation domain, PKS(KS) = PKS ketosynthase domain, CP = acyl or peptidyl carrier protein, TE = thioesterase.

We previously identified BGCs for actinonin and matlystatin^13^, which showed that these NPs are biosynthesised by either a non-ribosomal peptide synthetase (NRPS; actinonin^14^), or a hybrid NRPS/polyketide synthase (NRPS/PKS; matlystatin), where it was proposed that the HPS moiety was biosynthesised as a coenzyme A (CoA)-bound precursor and condensed at the beginning of the assembly line. A combination of bioinformatic analysis and stable isotope feeding led to a proposal for the biosynthesis of the HPS group (Fig. 1A), although the precise timing of biosynthetic steps remains to be determined. A key step in the proposed pathway is a mutase-catalysed rearrangement to generate the characteristic HPS carbon skeleton. This proposed mutase specificity towards hexylmalonyl-CoA would be specific to HPS biosynthesis, so we hypothesised that it could serve as an effective genetic marker for finding novel HPS-containing NPs.

*Rhodococcus fascians* is a plant pathogen that causes leafy gall disease across a broad range of plants and provides a significant economic burden for the ornamental plant industry^15–17^. *R. fascians* is the only plant pathogenic *Rhodococcus* species, where this pathogenicity is determined by the presence of a 199 kb linear virulence plasmid^18,19^ (pFiD188). Infection symptoms are a result of the production of *Rhodococcus* NPs that interfere with plant growth and development. *R. fascians* therefore represents an agriculturally important model for understanding how small molecules mediate microbial interactions in nature^20^. pFiD188 harbours multiple BGCs, including the *fas* BGC responsible for the biosynthesis of cytokinins, which induce the growth of differentiated plant tissue and can lead to gall formation^21^. The *att* BGC encodes for production of an autoregulatory compound proposed to be involved in the switch to an endophytic phase^22^, while the function of a pFiD188 NRPS BGC was unknown^18^.

Here, mutase-guided genome mining identified that the NRPS BGC on pFiD188 was predicted to make a putative metalloprotease inhibitor. We show that this BGC (*lyd*) produces the cyclic lipodepsipeptide lydiamycin A, which had been previously isolated from *Streptomyces* spp. as an antimycobacterial antibiotic^23–25^, but was not reported to have an HPS-like moiety. The synteny to the actinonin and matlystatin BGCs prompted a detailed revision of the lydiamycin structure to show that it features a rare HPS-like (*R*)-2-pentyl-succinyl chemophore. We show that lydiamycin is a PDF inhibitor via the identification of a self-immunity gene in the *lyd* BGC and demonstrate that lydiamycin production contributes to microbial competition during leaf colonisation.

## RESULTS

### Identification of putative metalloprotease inhibitor BGCs

To identify BGCs that may produce novel HPS-containing NPs, a phylogenetic analysis of all actinobacterial mutases was performed. A co-association analysis was then performed to determine whether homologues of proposed HPS biosynthesis genes and PKS/NRPS genes were present in the vicinity of mutase genes (Fig. S1), which revealed that a single clade was enriched with potential metalloprotease inhibitor BGCs (Fig. 1B, Table S5). This clade contained the known BGCs of actinonin (*Streptomyces* sp. ATCC 14903) and matlystatins (*Actinomadura atramentaria* DSM 43919), as well as uncharacterised BGCs from diverse genera, including *Actinoplanes, Micromonospora* and *Rhodococcus*.

*R. fascians* D188 was further investigated as it is a well-studied plant pathogen^16^. The BGC is located on the pFiD188 pathogenicity megaplasmid, which implicated the associated NP as a potential pathogenicity determinant, although the product was unknown^18^. This BGC (Figs. 2A and 3, Table S6), encodes multiple NRPS-associated proteins (LydDHJKL, Fig. S2), a crotonyl-CoA carboxylase-reductase (CCR, LydM), a hexylmalonyl-CoA epimerase (LydP), α and β subunits of a putative hexylmalonyl-CoA mutase (LydIN) and a MeaB-like protein (LydO), which could aid with mutase function^26^. The absence of genes homologous to the asparagine synthetase and *N*-oxygenase genes present in the actinonin and matlystatin BGCs suggested that the *R. fascians* product may instead feature a (*R*)-2-pentyl-succinyl chemophore (Fig. 1A) rather than the full hydroxamic acid moiety. The Stachelhaus specificity code^27,28^ of a standalone adenylation domain (LydK) is identical to the matlystatin NRPS MatJ (Fig. S2), which is predicted to activate piperazic acid^13,29^. The predicted incorporation of a piperazic acid residue was supported by the similarity of LydE and LydF to the functionally characterised KtzI/KtzT piperazic acid biosynthetic machinery described in kutzneride biosynthesis^30,31^ (47% and 45% identity, respectively). Taken together, these analyses suggested that this BGC makes a non-ribosomal peptide incorporating at least one piperazic acid residue and featuring an intermediate, carboxylated version of the chemophore.

**Figure 2.**
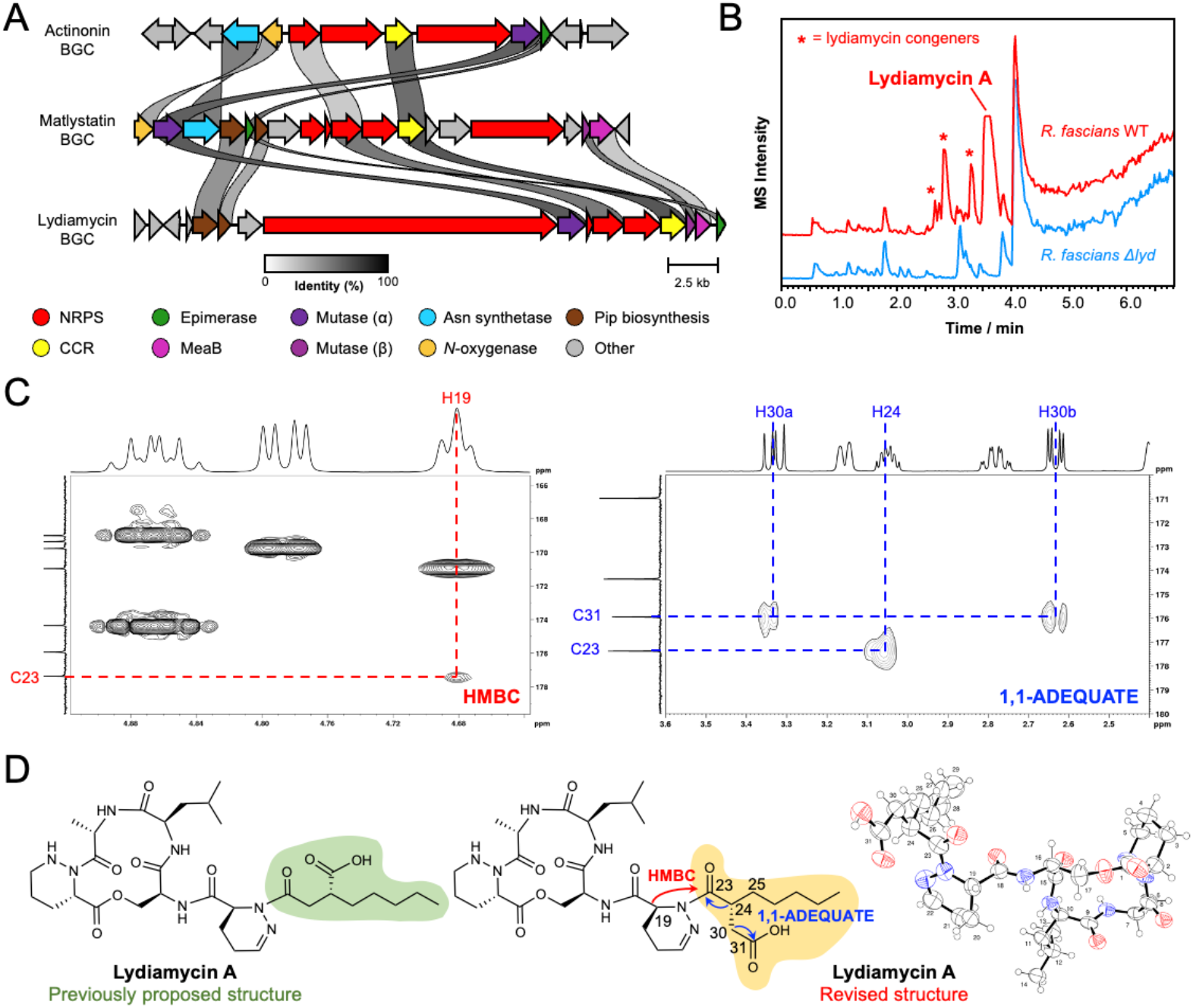
Characterisation of lydiamycin from *R. fascians*. A. Similarity comparison of the actinonin, matlystatin and lydiamycin BGCs. Gene identity percentage is represented by the grayscale links between genes. Pip = piperazic acid. B. Comparison of LC-MS chromatograms of WT R. fascians D188 and a strain with a deletion in the lydiamycin BGC (Δ*lyd*). C. Representative regions of 2D HMBC and 1,1-ADEQUATE NMR spectra with key correlations highlighted for the 2-pentyl-succinyl portion of the molecule. D. Comparison of the previously proposed lydiamycin A structure alongside the revised structure and the associated crystal structure.

**Figure 3.**
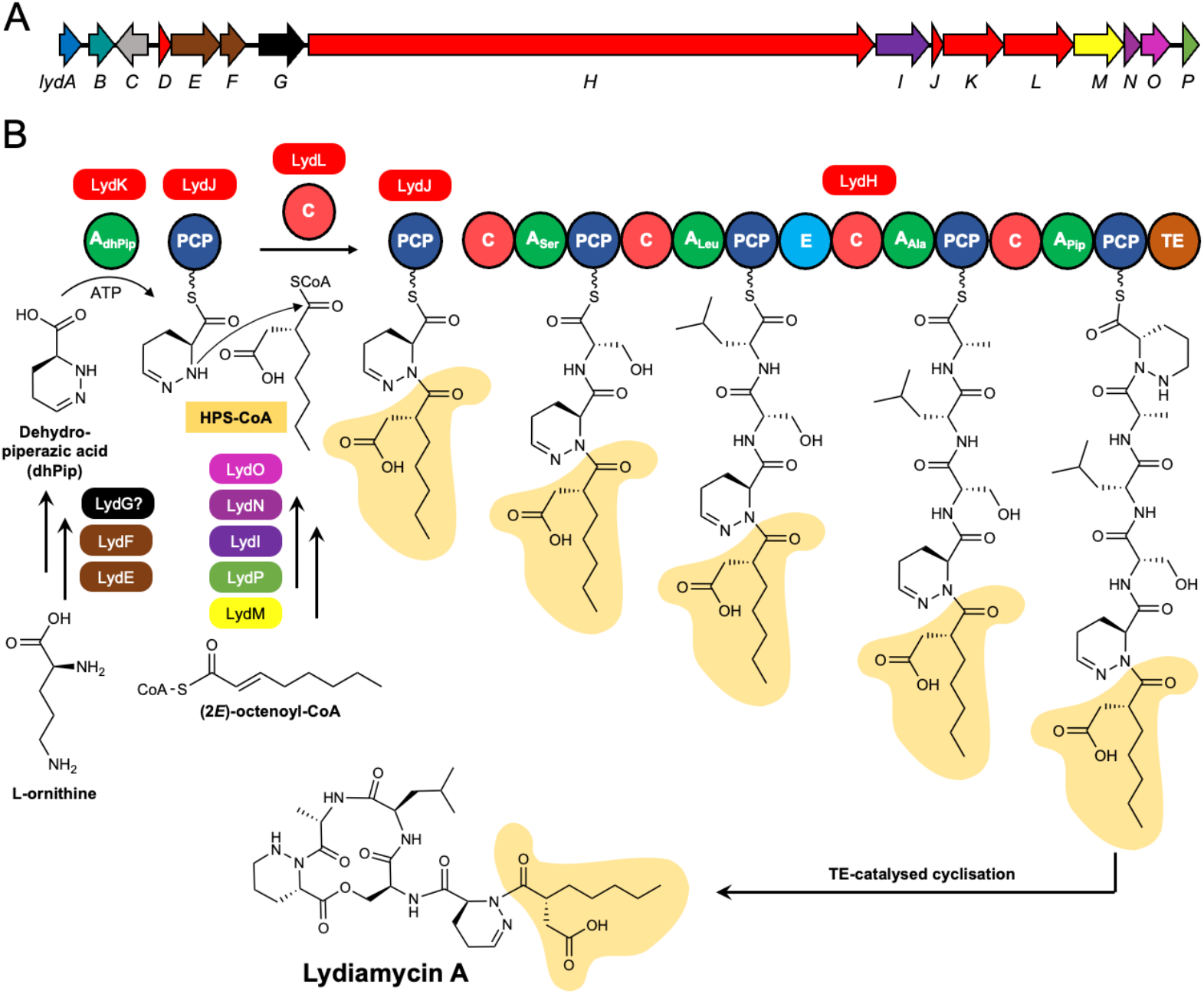
Proposed biosynthesis of lydiamycin A. A. *lyd* BGC is colour-coded as in Fig. 2A as well as genes encoding a PDF (LydA) in blue, a LuxR-type regulator in teal, a prenyltransferase in grey, a flavin-dependent oxidoreductase in black. See Table S6 for further BGC details. B. Biosynthetic proposal for lydiamycin. The timing of piperazic acid dehydrogenation is not known and may occur post-assembly line, given that LydK has an identical recognition sequence to Pip-selective MatJ in matlystatin biosynthesis (Fig. S2).

### The *R. fascians* D188 pathway makes lydiamycin A

To identify the product of the pFiD188 BGC, wild type (WT) *R. fascians* D188 and a mutant featuring a genetically disrupted adenylation domain^18^ (*R. fascians* D188 Δ*lyd*) were fermented in multiple media and sampled at multiple time points. The metabolomes were then compared using liquid chromatography-mass spectrometry (LC-MS), which provided 20 potential products of the BGC (Figs. 2B and S3-S5). Mass spectral networking analysis indicated that many of these compounds are likely to be structurally related (Figs. S6-S7). The mass of the major BGC-associated compound ([M+H]^+^ *m/z* 664.3680) (Fig. S8) was determined to be identical to lydiamycin A ([M+H]^+^ *m/z* 664.3665; 2.3 ppm). Further LC-MS/MS and nuclear magnetic resonance (NMR) spectroscopy analyses of this compound were also fully consistent with previously described lydiamycin A data^23,25^ (Fig. S9-S15, Tables S7-S9). These data indicate that the major product of the *R. fascians* D188 BGC is lydiamycin A and represents the first identification of the lydiamycin A BGC in a non-streptomycete. In parallel to our work, the first lydiamycin BGC was reported^32^ from a *Streptomyces* species, which is identical to the *Streptomyces aureoverticillatus* BGC in our phylogenetic analysis (Fig. 2A) and is very similar to the *R. fascians* lydiamycin BGC (Fig. S16).

Lydiamycins are cyclic depsipeptides first identified from *S. lydicus* HKI0343 and have been shown to inhibit mycobacteria, including activity towards a multidrug resistant clinical *M. tuberculosis* strain^23^. The published lydiamycin A structure is a macrocyclic pentapeptide with a pentyl-succinic tail^23–25^ (Fig. 2D) which partially matched our prediction that the D188 BGC product would feature piperazic acid and a terminal carboxylated moiety. The molecular networking data indicates that *R. fascians* D188 makes lydiamycin congeners that are distinct from the previously described lydiamycins A-H^23,33^.

### Bioinformatic Analysis Informs a Structure Revision of Lydiamycin A

A structural inconsistency of lydiamycin A is apparent when it is compared with actinonin and the matlystatins. Actinonin and matlystatin biosynthesis is proposed to proceed with the condensation of a peptide precursor with the carbonyl of the alpha-proximal thioester of a *N*-hydroxy-2-pentyl-succinamyl-CoA precursor^13^ (Fig.1A). However, the published lydiamycin A structure (Fig. 2D) necessitates the condensation of a dehydropiperazic acid residue to the carbonyl of the beta-proximal carboxylic acid of a 2-pentyl-succinyl precursor (Fig. S17). This disparity is not supported by the presence of any additional genes in the *lyd* BGC that could rationalise this alternative connectivity. Therefore, we hypothesised that lydiamycin A could be biosynthesised via the alpha-proximal condensation and thus instead features a HPS-like carbon skeleton consistent with actinonin and the matlystatins (Fig. 2D).

To test this proposal, a greater quantity of lydiamycin A was purified and analysed by NMR. A HMBC cross-peak with H19 allowed the distinction of the piperazic-proximal and piperazic-distal carbonyl carbon atoms as the δ_C_ 177.4 ppm and δ_C_ 176.0 ppm peaks, respectively (Fig. 2C). Consideration of the relative intensity of HMBC cross-peaks for the 2-pentyl-succinyl carbonyl carbon atoms was more supportive of the revised lydiamycin A structure than the published structure (Fig. S18). To further ratify this proposal, 1,1-ADEQUATE NMR (Fig. 2C, Fig. S19) showed that the piperazic-distal carbonyl carbon atom (C31) is adjacent to a carbon with a pair of diastereotopic protons (H30a, H30b; δ_H_ 3.33 and 2.63 ppm) and the piperazic-proximal carbonyl carbon atom (C23) is adjacent to a carbon with a single hydrogen (H24; δ_H_ 3.05 ppm). The proton multiplicity of these key carbon atoms was also confirmed by HSQC (Fig. S20). These observations are consistent only with the proposed revised lydiamycin A structure.

To determine the stereochemistry at C24, we reanalysed the phenylglycine methyl ester (PGME) derivatisation experiments published by Hwang and co-workers on lydiamycin from *Streptomyces* sp. GG23^25^. We applied the modified Mosher methodology described by Yabuuchi and Kusumi for using PGME derivatives to determine the absolute configuration of β,β-disubstituted propionic acids^34^. These data were fully consistent with *R* stereochemistry at C24 of lydiamycin (Fig. S21), which is also consistent with the biosynthetic proposal, as the chemophores found in actinonin and matlystatin are stereospecifically biosynthesised as *R* enantiomers^5,35^. To further support this structural revision, we acquired X-ray crystallography data from ∼30 µm^3^ crystals of lydiamycin A, which confirmed the structure at a 0.8 Å resolution (Fig. 2D).

### Lydiamycin A inhibits peptide deformylase

Actinonin is a potent inhibitor of peptide deformylase (PDF)^6^, whereas the mechanism of action of lydiamycin had not been determined and the structural similarity to actinonin had previously not been noted. We therefore hypothesised that a PDF gene at the edge of the *lyd* BGC (*lydA*) could represent a self-immunity mechanism and that PDF is the molecular target of lydiamycin. Lydiamycin lacks the hydroxamate functionality that enables actinonin and matlystatin to tightly bind active site metals, but the carboxylate functionality of the (*R*)-2-pentyl-succinyl moiety could potentially still bind into the PDF active site.

Since lydiamycin was previously shown to display anti-mycobacterial activity^23^, the non-pathogenic model organism, *Mycobacterium smegmatis* mc^2^155 was used to assess the role of *lydA in vivo*. Exogenous overexpression of *lydA* rescued *M. smegmatis* to wild-type-like growth when exposed to 50 μM lydiamycin A (Fig. 4A), which was sufficient to prevent growth in the empty vector control or in a strain of *M. smegmatis* overexpressing its native housekeeping PDF gene (Fig. S22). This suggests that the *lydA*-mediated resistance is not caused by a titration of lydiamycin A but rather an inherent resistance to the compound. Given that the *lyd* BGC is encoded on plasmid pFiD188, we assessed the lydiamycin sensitivity of wild type *R. fascians* D188 and a pFiD188-free strain (*R. fascians* D188-5). *R. fascians* D188-5 was determined to be at least 60 times more sensitive to lydiamycin A than the *R. fascians* wild type strain (MIC = 10 ug/mL versus >600 μg/mL, respectively; Fig. S23). These data indicate that LydA functions as a self-immunity determinant.

**Figure 4.**
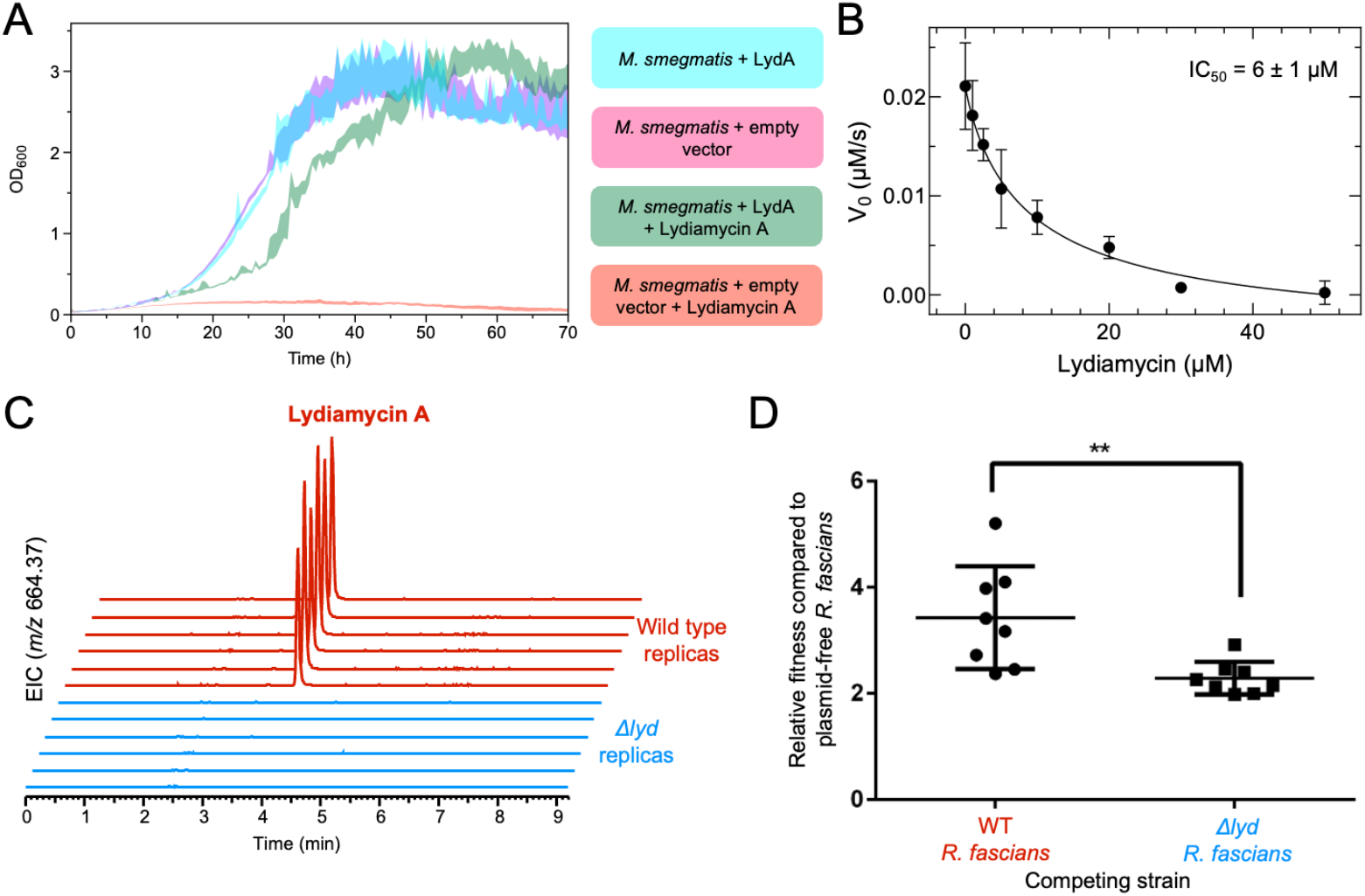
Biological activity of lydiamycin A. A. Growth curves of *M. smegmatis* expressing either empty vector or the PDF gene encoded in the lydiamycin BGC (LydA), in the absence or presence of 50 μM lydiamycin A. The standard error of the mean is illustrated by the shaded envelope (n=3 for controls, n=6 for experiments with lydiamycin). B. *In vitro* inhibition of *E. coli* PDF by lydiamycin A. Initial velocities V0 of PDF activity with a model peptide (formyl-methionyl-leucyl-p-nitroaniline, Fig. S24) were fitted to a sigmoidal function. Error bars represent the standard deviation of three independent experiments, n=3. C. Production of lydiamycin A by *R. fascians* during colonisation of *N. benthamiana* seedlings. Extracted ion chromatograms (EICs) are shown for *m/z* 664.3665 (+/-5 ppm) from LC-MS experiments. D. Competition assay of plasmid-free *R. fascians* (D188-5) against WT and Δ*lyd* strains. The relative fitness of each test strain was calculated as compared to the plasmid-free strain. Individual replicates are indicated as dots. The mean of these replicates is shown as a black cross-bar and error bars indicate the standard error of the mean. A two-tailed t-test showed a significant different between WT and Δ*lyd* relative fitness values (*p* = 0.0068).

To understand whether lydiamycin A directly inhibits PDF, *in vitro* assays were conducted with *Escherichia coli* PDF^36,37^, which showed that lydiamycin A exhibits dose-dependent inhibition of this PDF with an IC_50_ of 6 μM (Fig. 4B and S24). There is precedence for an intermediate carboxylated group to confer PDF inhibitory bioactivity, as the same moiety is present in the PDF inhibitor Sch-382583 isolated from a *Streptomyces* species^38^, while carboxylated variants of actinonin and matlystatin can inhibit PDF, albeit with significantly reduced potencies^39^. These data indicate that lydiamycin is an antimycobacterial agent that functions by peptide deformylase inhibition. There are no PDF genes in the actinonin or matlystatin BGCs, which means that *lydA* represents a novel self-immunity gene associated with the production of a PDF inhibitor. Our genome mining analysis identified multiple BGCs that do encode PDFs (Fig. 1B), indicating that this strategy may exist in currently uncharacterised pathways.

### Lydiamycin A is an important ecological agent during niche colonisation

As *R. fascians* is the causative agent of leafy gall disease, lydiamycin A was investigated for its involvement in leafy gall disease pathogenesis in *Nicotiana benthamiana* plants^19^, especially as actinonin had previously been identified as a chloroplast PDF inhibitor^40^. However, there was no significant difference between plant or leafy gall mass for plants infected with either wild type *R. fascians* or *R. fascians* Δ*lyd* and grown for 4 weeks (Fig. S25). Equivalent results were obtained for assays for the root length of seedlings and gall development on excised leaves (Figs. S26-S27). These data indicate that the production of lydiamycin A is not required for leafy gall disease development and does not have phytotoxic activity. However, LC-MS analysis of *Nicotiana* seedlings and excised leaves infected with either wild type *R. fascians* or *R. fascians* Δ*lyd* (Fig. 4C) indicates that lydiamycin A is reliably produced *in planta* during *R. fascians* infection and suggests that lydiamycin may have an ecologically important role.

We hypothesised that the production of lydiamycin during leaf colonisation could instead reflect a role in microbial competition, which was tested by conducting competition assays with plasmid-free *R. fascians* D188-5, which is sensitive to lydiamycin (Fig. S23). Here, the impact of lydiamycin production was determined by inoculating sterile *N. benthamiana* seedlings with *R. fascians* D188 WT or Δ*lyd* in a 1:1 mixture with plasmid-free (and lydiamycin sensitive) D188-5. After one week of plant growth, bacterial cells were recovered and the relative amounts of each strain were determined. This experiment indicated that lydiamycin production provides a significant fitness benefit based on the relative fitness ratio of WT/D188-5 compared to Δ*lyd*/D188-5 (Fig. 4D), which is consistent with production of an antibiotic inhibiting the growth of a sensitive competitor. More generally, carriage of pFiD188 provides a significant fitness benefit to *R. fascians* during plant colonisation given that D188-5 was outcompeted in both competition assays. A repeat experiment showed a comparable competitive advantage (Fig. S28). These data suggest that lydiamycin A production may be important for antagonising rival bacteria that compete for similar resources and therefore enhances the likelihood of *R. fascians* successfully colonising plants.

## DISCUSSION

Metalloproteinase inhibitors are promising lead molecules for a variety of therapeutic targets^41^. The HPS moiety in actinonin represents a potent chemophore for the inhibition of multiple therapeutically relevant proteases and hydrolases, including PDF^42^. However, currently few NPs have been discovered with a similar chemophore. Here, mutase-guided genome mining enabled us to identify multiple candidate BGCs that we hypothesised produce protease inhibitors. Our discovery that a BGC in *R. fascians* makes lydiamycin was initially unexpected given the dissimilarity between the published lydiamycin structure and actinonin, but extensive NMR analysis, reanalysis of published PGME derivatisation data^25^ and X-ray crystallography fully supported the structural revision of lydiamycin A (Fig. 2D) to show that it features a (*R*)-2-pentyl-succinyl moiety with the same carbon connectivity and stereochemistry as HPS in actinonin.

This discovery highlights how consideration of the biosynthetic pathway can be an important tool to inform structural nuances that could be overlooked by chemical derivatisation or routine NMR analysis^43^, which has previously been demonstrated with the biomimetic synthesis and subsequent structural revision of NPs such as hyperelodione D^44^. Multiple previous synthetic and analytical studies had been unable to determine the true connectivity of lydiamycin^23,24,33^, where each analysis lacked the critical HMBC correlation between C23 and H19. The connectivity at this part of the molecule was instead originally defined via MS/MS fragmentation^23^. A full understanding of the structure and stereochemistry of lydiamycin A now enables accurate structure-activity relationship experiments to be undertaken for both the peptide portion and the (*R*)-2-pentyl-succinyl moiety, especially as its affinity to metalloproteinases is predicted to be modulated by changing the carboxylate to a hydroxamate^39^. Metabolomics-based identification of multiple novel lydiamycin congeners from *R. fascians* (Figs. S3-S7) provides further natural structural diversity to test for activity, as the masses of these molecules differ to lydiamycin congeners previously identified in other producing organisms^23,33^. Notably, the medium had a significant effect on the array of lydiamycin-like molecules produced. These observations highlight the value of investigating related BGCs in different producers, as they may have the potential to make chemodiverse congeners for SAR studies and pharmacological optimisation. For example, novel congeners of the lipodepsipeptide ramoplanin were obtained by the identification of related BGCs^45^.

*R. fascians* produces the LydA PDF to confer immunity against lydiamycin. There are several reported methods for bacteria gaining resistance against PDF inhibitors, including by PDF mutations^46^, maintaining protein synthesis by bypassing formylation^47,48^, efflux of the inhibitor^49^ or PDF overexpression^50^. Overexpression of the housekeeping *M. smegmatis* PDF gene did not confer resistance, which infers that LydA is truly resistant to lydiamycin. Further biochemical work is required to understand the precise structural determinants of this LydA immunity, especially as none of the mutations reported in the literature that confer PDF resistance to inhibitors are present in LydA^46,51^, suggesting that it confers resistance in a novel manner. Our identification of PDF genes in multiple BGCs indicates that this mechanism of immunity may exist for other PDF inhibitors and that PDF genes may represent underexplored markers of antibiotic BGCs, as highlighted by the recent discovery of gammanonin^52^. Co-association of BGCs with copies of antibiotic target genes represents a promising method for antibiotic discovery and an associated understanding of mechanism of action^53–55^, which can be aided by informatic tools such as ARTS^56^.

PDFs constitute highly promising antimicrobial targets, given the fitness trade-off observed for acquired resistance phenotypes^48^. However, hydroxamate-containing PDF inhibitors, such as actinonin, have potentially not succeeded in clinical trials due to their toxicity, which may be associated with the *in vivo* conversion of this chemical moiety into mutagenic isocyanates via Lossen rearrangement^57^, which has been implicated in the mutagenicity of hydroxamate-based histone deacetylase inhibitors (HDACIs)^58^. Our work highlights how multi-layered genome mining can select for BGCs producing PDF inhibitors carrying the same chemophore backbone but armed with a non-toxic carboxylate as metal chelator. It is possible that natural PDF inhibitors have similar functional logic as synthetic hydroxamate-based HDACIs. These compounds are normally composed of three parts: a cap group that interacts with the surface of the enzymatic target, a linker and a metal chelator moiety^58^. Given that matlystatins and actinonin share the same HPS chemophore, it is conceivable that the rest of the molecules acts as the specificity determinant cap. Our definition of the lydiamycin NRPS machinery opens the possibility of creating optimized metalloproteinase inhibitors by combining natural specificity-determinant caps and non-toxic carboxylic acid chelators through combinatorial biosynthesis and NRPS engineering^59^.

*R. fascians* is an increasingly economically relevant plant pathogen, where it can form long-term biotrophic interactions with the plant host and triggers development of differentiated leafy galls and fasciation^15,16^. The *lyd* BGC is present on the pFiD188 pathogenicity megaplasmid of *R. fascians*, although we found no evidence for lydiamycin involvement in pathogenesis, despite the sensitivity of *Nicotiana* chloroplast PDF to actinonin^40^. Instead, leaf-based competition experiments indicate that lydiamycin may play a role in niche colonisation (Fig. 4D). This activity may indirectly impact the establishment of disease and long-term plant colonisation via competition within a complex plant microbiome. Bacterial communities associated with plants are known to be a rich source of antibiotics^60–62^, where they can function to control plant disease^63,64^. The production of antibiotics by pathogens themselves is less well understood or investigated, despite the ability of multiple pathogens to make potent antibiotics^65–67^. An alternative model for lydiamycin production could relate to plasmid fitness, where the capacity to make an antibiotic and confer resistance to the antibiotic could feasibly help counteract the fitness cost of plasmid carriage^68^. Production of a diffusible antibiotic could also deter plasmid-free cheater *Rhodococcus* strains in a mixed population, as it is known that plasmid-free *R. fascians* is capable of effectively colonising plants^20,69,70^. These models should be investigated in future experiments with *R. fascians*.

In conclusion, our identification of the lydiamycin BGC on the *R. fascians* pFiD188 pathogenicity plasmid has guided the structural recharacterisation of lydiamycin, the identification of its molecular target and prompted an investigation of its role for plant colonisation. Our work should stimulate the rational discovery of further PDF inhibitors, as well as highlighting the potential ecological functions of antibiotic production by microbes.

## ACKNOWLEDGEMENTS

This work was funded by a UK Research and Innovation Biotechnology and Biological Sciences Research Council (BBSRC) Norwich Research Park Doctoral Training Partnership grant (BB/M011216/1) for J.J.F. The work was also supported by BBSRC Institute Strategic Programme grants (BBS/E/J/000PR9790 and BB/X01097X/1) for the John Innes Centre (JIC). J.W.S. was funded by BBSRC grant BB/T015349/1. The authors wish to acknowledge the Diamond Light Source for access to beamline i04 under proposal MX32728. This work was supported by the excellent technical assistance at JIC provided by Dr Lionel Hill and Dr Carlo Martins for LC-MS, Dr Martin Rejzek for chemical purification, and Prof. David Lawson and Julia Mundy for crystallography. We are thankful to Dr Catriona Thompson (JIC) for assistance with the statistical analysis of data, and to Prof. Barrie Wilkinson (JIC) for helpful discussions relating to the project.

