## Supporting Information for "Discovery of lydiamycin A biosynthetic gene cluster in the plant pathogen *Rhodococcus fascians* guides structural revision and identification of molecular target"

### MATERIALS AND METHODS

#### Chemicals and reagents

All chemicals and media components were purchased from Sigma Aldrich, except for the following: agar (Melford), NaCl and glucose (Fisher Scientific), yeast extract (Merck), soya flour (Holland and Barrett) and peptone (BD Biosciences). All enzymes were supplied by New England Biolabs (NEB) unless otherwise specified. All solvents for extractions and chromatography were supplied by Fisher Scientific. Ultrapure water was obtained using a Milli-Q purification system (Merck). All media were autoclaved prior to use and chemical solutions were filter sterilised using a 0.22 μM syringe filter. All media used are defined in Table S2 and were made up in 1 L of Milli-Q water. Solid media were made with 2% agar unless stated otherwise. Antibiotics were added where necessary to the following concentrations: kanamycin (50 μg/mL for *E. coli* and 20 μg/mL for *M. smegmatis*); chloramphenicol (25 μg/mL), streptomycin (100 μg/mL). All primers were synthesised by Eurofins Genomics or Integrated DNA Technologies at a 25 nmol synthetic scale with standard desalting. Primers were diluted to 100 μM in Milli-Q water and stored at -20 °C. Working stocks were prepared at 10 μM and stored at 4 °C.

#### General microbiology methods

Unless otherwise stated, *R. fascians* strains were grown in liquid LB medium at 28 °C with shaking at 250 rpm for 24-48 hours or on solid LB medium at 28 °C until colonies were visible (2-3 days). Unless otherwise stated, *M. smegmatis* cultures were grown aerobically in liquid MOADC medium at 37 °C with shaking at 250 rpm for 18-24 hours or on solid MOADC for 3 days. Sterile glass beads and 0.25% TWEEN-80 were added to liquid cultures to reduce cellular aggregation. Unless otherwise stated, *E. coli* strains were grown in liquid LB medium at 37 °C with shaking at 250 rpm for 16-18 hours or on solid LB medium at 37 °C. Plates of all strains were stored at 4 °C. Long-term stocks of all strains were stored at -70 °C in 25% glycerol. gDNA was extracted from *R. fascians* and

*M. smegmatis* using the Fast DNA SPIN Kit for Soil (MP Biomedicals), according to the manufacturer's protocol. The concentration and purity of gDNA, purified amplified DNA and digested vector backbones was determined using a NanoDrop 2000 spectrophotometer (Thermo Scientific) according to the manufacturer's protocol.

**Identification and phylogeny of actinobacterial acyl-CoA mutases**

The MatB protein sequence (WP_019634562.1) was used in an InterPro[1] search to identify the methylmalonyl-CoA mutase InterPro entry (IPR006099). All actinobacterial protein sequences belonging to the IPR006099 InterPro entry (~13,000 sequences at time of analysis, February 2022) were downloaded in .FASTA format. Mutase protein sequences identified in a MatB BLAST[2] analysis that were highly similar (>60% identity) but absent in the InterPro database were added to the sequence list (WP_019634562.1, WP_176725504.1, WP_210837750.1, WP_063760750.1, WP_189742219.1, WP_223771548.1, WP_104814753.1, WP_184992507.1, QES31823.1, QNF54062.1, QHF95772.1). Sequences were submitted to the Enzyme Similarity Tool (Enzyme Function Initiative; EFI-EST[3]) to produce a protein sequence similarity network (SSN). Edges were drawn between nodes where the similarity e-value was less than $e^{-50}$. Protein sequences of >40% sequence similarity were collated into 1,615 representative nodes to generate the SSN. The SSN was exported and visualised using Cytoscape[4]. The SSN node table was exported as a .csv file and the UniProt retrieval tool was used to obtain the protein sequences of the first accession number of each representative node ("*representative FASTA file*"). Representative accessions are shown in Table S5. The resulting protein sequences were then aligned using ClustalW[5] with default parameters at the CIPRES server[6]. A tree was then constructed with this alignment file using the 'RAxML-HPC Blackbox' method[7] with default parameters at the CIPRES server, which determined that a bootstrap value of n=350 was sufficient. The tree was then visualised using iTOL[8].

**Co-association analysis of mutases and other genes**

In order to perform a co-association analysis with one protein sequence from each representative node of the SSN described above, the *representative FASTA file* was submitted to EFI-EST[3] to generate a network lacking clustering (ID=100%). This SSN was then submitted to the Enzyme Function Initiative Genome Neighbourhood Tool (EFI-GNT)[3]. Parameters of neighbourhood size = 15 and minimal co-occurrence percentage lower limit = 20 were set. The resultant SSN was viewed in Cytoscape[4] and the node table exported as a .csv file, which was then assessed for Pfam domains of interest for each accession number. The Pfam domains of interest were as follows: PF00107 (CCR), PF13669 (acyl-CoA epimerase), PF02310 (acyl-CoA mutase β subunit), PF03308 (MeaB-like), PF00733 (asparagine synthetase), PF11583 (*N*-oxygenase), PF02770 (acyl-CoA dehydrogenase), PF00501 (NRPS, adenylation), PF00668 (NRPS, condensation), PF00109 (PKS, ketosynthase), PF00550 (carrier protein) and PF00975 (thioesterase). Co-associated protein domains were mapped to the phylogenetic tree using iTOL[8].

**Phylogeny and co-association analysis of the BGC clade**

A clade was identified in the above analysis that co-associated with multiple biosynthetic Pfam domains. The representative nodes of this clade were therefore subjected to another phylogenetic and co-association analysis. All protein sequences from each representative node were retrieved from the original EFI-EST SSN, in addition to a phylogenetic outgroup (UniProt accession A0A166QLU4). The >60% identity sequences absent in the UniProt database were also added, as detailed above. In total, 45 sequences were submitted to CIPRES for ClustalW alignment and RAxML-HPC Blackbox tree building, using default parameters. The program determined that a bootstrap value of n=300 was sufficient and de-replicated 20 sequences that were 100% identical. The co-association analysis was performed using the same parameters as detailed above, along with additional Pfam domains of interest: PF13434 (KtzI-like), PF04299 (KtzT-like), PF07690 (transporter) and PF01327 (peptide deformylase). This co-association analysis was converted into a binary annotation file and displayed on the tree using iTOL. The tree was manually trimmed to remove phylogenetically identical Acyl-CoA mutase leaves that had identical co-association results.

**Bioinformatic Analysis of the lydiamycin BGC**

antiSMASH 6.2[9] was used for general analysis of the lydiamycin BGC. NRPSsp[10], PKS/NRPS Analysis[11] and NRPSpredictor2[12] were used for predicting the adenylation specificity of the *lyd* NRPS genes. Stachelhaus sequences were obtained from the antiSMASH output. clinker[13] was used to compare BGCs. Lydiamycin BGC data for the *S. venezuelae* (NZ_CP029193.1) and *S. aureoverticillatus* (OL452061.1) clusters was used.

**Initial screening of *R. fascians* metabolite production**

*Rhodococcus fascians* D188 and *R. fascians* D188 Δ*lyd* were grown in liquid LB (Table S2) at 28 °C, 250 rpm under aerobic conditions. *R. fascians* D188 Δ*lyd* was first described in Francis *et al.*[14] as Δ*nrp5.* Pre-cultures were grown for 2 days and 1% used for inoculations. The following liquid media were selected for testing *R. fascians* metabolite production and are described in Table S2: aurachin production medium[15], rhodostreptomycin production medium[16], minimal media A (MinA)[17], screening medium 7 (SM7), SM12, yeast extract broth (YEB), actinonin production medium[18], matlystatin production medium[18], and bottromycin production medium[19]. In the comparative metabolomics experiment, 10 mL of each of the above media, in sterile bunged 50 mL falcon tubes, were inoculated with 1% *R. fascians* D188 WT and *R. fascians* D188 Δ*lyd* pre-cultures in duplicate. Samples were incubated at 28 °C, 250 rpm and sampled at 5 and 12 days by removing 0.75 mL of culture. This was extracted in 0.75 mL MeOH (shaken, 1 hour), centrifuged (13,000 rpm, 30 minutes) and 0.6 mL of supernatant aliquoted for LC-MS analysis. Medium-only samples were prepared using the same method.

Samples were subjected to LC-MS analysis using a Shimadzu Nexera X2 UHPLC coupled to a Shimadzu IT-ToF mass spectrometer. Samples (5 µL) were injected onto a Phenomenex Kinetex 2.6 µm C18 column (50 x 2.1 mm, 100 Å), eluting with a linear gradient of 5 to 100% acetonitrile in water + 0.1% formic acid (FA) over 6 minutes with a flow-rate of 0.6 mL/min at 40 °C. Positive mode mass spectrometry data were collected between *m/z* 300 and 2000 with an ion accumulation time of 10 ms featuring an automatic sensitivity control of 70% of the base peak. The curved desolvation line temperature was 300 °C and the heat block temperature was 250 °C. MS/MS data was collected in a data-dependent manner using collision-induced dissociation energy of 50% and a precursor ion width of 3 Da. The instrument was calibrated using sodium trifluoroacetate cluster ions prior to every run. LabSolutions software (Shimadzu) was used to identify molecules present in WT *R. fascians* and absent in *R. fascians* Δ*lyd*.

**Mass spectral networking**

Single colonies of *R. fascians* D188 WT and Δ*lyd* strains were used to inoculate 10 mL LB and grown for 2 days at 30 °C. These cultures were used to inoculate (1% v/v) 10 mL YEB and SM12 media in in triplicate in 50 mL sterile bunged falcon tubes. Media only samples were also prepared in triplicate. All cultures were incubated at 30 °C with shaking at 250 rpm for 5 days. 0.75 mL aliquots of the cultures were extracted in 0.75 mL MeOH for 1 hour, with shaking. Samples were centrifuged

(15,871 x *g*) for 30 minutes. The resulting supernatants were subjected to LC-MS/MS analysis using a Waters Acquity UHPLC coupled to a Q-Exactive Orbitrap Mass Spectrometer (Thermo). Samples (5 µL) were injected onto a Phenomenex Kinetex 2.6 µm C18 column (50 x 2.1 mm, 100 Å), eluting with a linear gradient of 5 to 95% acetonitrile in water + 0.1% FA over 6 minutes with a flow rate of 0.6 mL/min at 40 °C. Positive mode mass spectrometry data was collected between *m/z* 150 and 2000. MS/MS data were collected in a data-dependent manner with the following parameters: chromatography peak width = 7 s; resolution = 70,000, AGC target = $3x10^6$ , maximum IT = 100 ms, scan range 150 to 2000 *m/z*; dd-MS2 settings: resolution 17,500, AGC target = $1x10^5$ , maximum IT = 50 ms, loop count = 5, isolation window 1.5 *m/z*, isolation offset = 0.0 *m/z*, stepped Normalized Collision Energy = 20, 40, 60; dd settings: minimum AGC target = $8x10^3$, Intensity threshold = 1.6 $x10^5$, exclude isotopes = ON, dynamic exclusion = 1 s.

Data was converted to mzML format and molecular networks were created using the online workflow on the GNPS website[20,21]. The data was filtered by removing all MS/MS fragment ions within +/- 17 Da of the precursor m/z. MS/MS spectra were window filtered by choosing only the top 6 fragment ions in the +/- 50 Da window throughout the spectrum. The precursor ion mass tolerance was set to 0.1 Da and a MS/MS fragment ion tolerance of 0.1 Da. A network was then created where edges were filtered to have a cosine score above 0.7 and more than 6 matched peaks. Further, edges between two nodes were kept in the network if and only if each of the nodes appeared in each other's respective top 10 most similar nodes. Finally, the maximum size of a molecular family was set to 100, and the lowest scoring edges were removed from molecular families until the molecular family size was below this threshold. The spectra in the network were then searched against GNPS spectral libraries. The library spectra were filtered in the same manner as the input data. All matches kept between network spectra and library spectra were required to have a score above 0.7 and at least 6 matched peaks. The network was visualised using Cytoscape (version 3.8.2). The GNPS results are available at:
https://gnps.ucsd.edu/ProteoSAFe/status.jsp?task=648c5adcb06b4733ba6ef092bcad9153

**Lydiamycin A purification**
10 mL of LB was inoculated with a single colony of WT *R. fascians* D188 and grown for 2 days at 30 °C. 0.5 mL of this culture was used to inoculate 50 mL of LB in a 250 mL flask, which was cultured at 30 °C for 1 day with 250 rpm shaking. This pre-culture was used to inoculate 2 x 1 L of SM12 medium (in 2000 mL flasks) at 1% v/v. These cultures were fermented at 30 °C for five days with shaking at 250 rpm. The cultures were centrifuged at 12,000 x *g* for 30 minutes and the resulting supernatant was extracted in two volumes of ethyl acetate for an hour. The organic extract was washed with an equal volume of water three times. The organic extract was dried over $MgSO_4$ and then dried *in vacuo* to yield 140 mg of organic extract. This material was resuspended in 3 mL MeOH and loaded onto a 30 g Sfär C18 cartridge (Biotage), pre-equilibrated with 5% acetonitrile in water. The sample was fractionated by flash chromatography (Biotage Isolera) at 12 mL/min with UV monitoring at 210 and 237 nm. Mobile phase A: water; mobile phase B: acetonitrile; elution started from 5% B with a gradient to 100% B over 10 column volumes (CV), and then holding at 100% B for 3 CV. The fractions were analysed by LC-MS and those containing lydiamycin A were combined, dried *in vacuo* and further purified by preparative HPLC using a Dionex UltiMate 3000 HPLC instrument (Thermo Scientific). To do this, the sample was dissolved in MeOH to 9 mg/mL, filtered using a 0.45 µm PTFE filter (Whatman) and applied to a Gemini-NX C18 (150 mm x 21.1 mm) column (Phenomenex). The mobile phase used was A: water with 0.1% formic acid; mobile phase B: acetonitrile with 0.1% formic acid; flow rate 20 mL/min; injection volume 500 µL; elution gradient:

$T$ = 0 min, 5% B; $T$ = 2 min, 5% B; $T$ = 7 min, 45% B; $T$ = 23 min, 70% B; $T$ = 25 min, 95% B; $T$ = 90 min, 95% B; $T$ = 30.1 min, 5% B; $T$ = 33 min, 5% B; UV absorbance monitoring at 237 nm. The major peak was collected and dried *in vacuo* to afford 20 mg of lydiamycin A as a white powder.

**Lydiamycin A structural elucidation**

High-resolution mass spectra were acquired on a Synapt G2-Si mass spectrometer equipped with an Acquity UPLC (Waters). Aliquots of the samples were injected onto an Acquity UPLC® BEH C18 column, 1.7 µm, 1x100 mm (Waters) and eluted with a gradient of acetonitrile/0.1% FA (B) in water/0.1% FA (A) with a flow rate of 0.08 mL/min at 45 °C. The concentration of B was kept at 1% for 1 min followed by a gradient up to 40% B in 9 min, ramping to 99% B in 1 min, kept at 99% B for 2 min and re-equilibrated at 1% B for 4 min. MS data were collected in positive mode with the following parameters: resolution mode, positive ion mode, scan time 0.5 s, mass range *m/z* 50-1200 calibrated with sodium formate, capillary voltage = 2.5 kV; cone voltage = 40 V; source temperature = 125 °C; desolvation temperature = 300 °C. Leu-enkephalin peptide was used to generate a lock-mass calibration with 556.2766, measured every 30 s during the run. For MS/MS fragmentation, a data directed analysis (DDA) method was used with the following parameters: precursor selected from the 4 most intense ions; MS2 threshold: 5,000; scan time 0.5 s; no dynamic exclusion. In positive mode, collision energy (CE) was ramped between 10-30 at low mass (*m/z* 50) and 15-60 at high mass (*m/z* 1200).

For NMR acquisition, pure lydiamycin A (7.0 mg) was dissolved in 200 µL $CDCl_3$ and transferred into a 3 mm NMR tube. This sample was subjected to a series of 1D and 2D NMR experiments on a Bruker Avance Neo 600 MHz spectrometer equipped with a TCI cryoprobe at 298 K. The NMR experiments carried out were Proton (16 scans), Carbon (900 scans), HMBC (8 scans), COSY (1 scan), HSQC (6 scans), 1,1-ADEQUATE (197 scans). Spectra were analysed using Bruker TopSpin 4.0.

**Lydiamycin A crystal structure**

A concentrated solution of lydiamycin A was prepared in toluene with gentle heating then DMSO was dropwise close to insolubility. The flask was then sealed and stored at 4 °C and assessed for crystal formation. Suitable crystals were resuspended in 100% ethylene glycol and harvested using Litholoops (Molecular Dimensions), then flash-cooled by plunging into liquid nitrogen prior to transport to the synchrotron. X-ray data were recorded on beamline I04 at the Diamond Light Source (Oxfordshire, UK) using an Eiger2 16M detector (Dectris) with the crystal maintained at 100 K by a Cryojet cryocooler (Oxford Instruments). The crystals diffracted very strongly, but at an X-ray wavelength of 0.6888 Å, the maximum resolution of the inscribed circle at the minimum possible detector distance was only 0.95 Å. In order to obtain complete data beyond this resolution, diffraction data were taken from two separate crystals, and for each of these, 4 x 360° passes were recorded at increasing $\chi$ offsets of 0, 15, 30 and 45°. Each pass comprised 3600 x 0.1° images with an estimated total dose of 0.5 MGy.

The X-ray data were integrated and scaled using DIALS[22]. Cell parameters for each sweep were refined independently using all strong indexed reflections. All sweeps were combined into a single file and reflections were scaled but not merged. An absorption correction was applied using the Blessing model and a median unit cell, calculated from the individual sweeps, was applied to the output file. Data beyond 0.8 Å resolution were excluded from the subsequent analysis. The space group was *C*2 with cell parameters of $a$ = 20.9, $b$ = 14.7, $c$ = 24.2 Å, $\beta$ = 97.3°. The structure was solved using SHELXT-2018[23] and refined on $F^2$ using SHELXL-2018[24]. Two molecules of lydiamycin A were present in the asymmetric unit. In both molecules, the carbonyl oxygen (O(1)) of the ester linkage was disordered over two positions. In one of the molecules, the terminal CH3CH2 fragment of the pentyl side-chain (C(28) and C(29)) was disordered over two positions. After completion of the model for lydiamycin A, voids remained in the structure. A solvent mask was calculated in Olex2[25], and 262 electrons were found in a volume of 822 $Å^3$ in 2 voids per unit cell. This is consistent with the presence of six water molecules per asymmetric unit which account for 240 electrons per unit cell. At the completion of refinement, the values of the Flack and Hooft parameters were ambiguous: 0.97(18) and –0.2(2), respectively. Absolute configuration was therefore assigned by consideration of the known centres C(2), C(7), C(10), C(16) and C(19), and the NMR data for assignment of C(24). At the conclusion of refinement, $R^1$ = 6.45%, $wR^2$ = 19.73%, GOOF = 1.067. Data were deposited with the Cambridge Crystallographic Data Centre with reference 2377371.

**Generation of pJAM vectors for PDF gene expression in *M. smegmatis***

Genes encoding the housekeeping *M. smegmatis* PDF (*MsPDF*, WP_011727226.1) and the PDF from the *R. fascians* lydiamycin BGC (*lydA*) were amplified from gDNA using Q5 polymerase (NEB) and a T100 Thermal Cycler (Bio-Rad). PCR reactions were separated by gel electrophoresis and purified using the GFX PCR DNA and Gel Band Purification Kit (Cytiva) according to the manufacturer's protocol. The *lydA* gene and pJAM2 vector[26] were digested with BamHI and XbaI and then gel purified. *lydA* was ligated into pJAM2 using T4 DNA ligase (Thermo) and the reaction was used to transform chemically competent *E. coli* DH5α cells. *MsPDF* was inserted into pJAM2 using NEBuilder HiFi DNA Assembly Master Mix (NEB). The reaction was incubated at 50 °C for 60 minutes and 2 µL of the reaction mixture used to transform chemically competent *E. coli* DH5α cells. All vectors were sequenced by Eurofins Genomics using a Mix2Seq Kit.

**Genetic manipulation of *M. smegmatis***

10 mL of MOADC (Table S2) was inoculated with a single colony of *M. smegmatis* and grown overnight at 37 °C. 100 mL of MOADC (in 250 mL flask) was inoculated with the pre-culture (1% v/v) and grown at 37 °C until an $OD_{600}$ of 0.2-0.8 was reached. The culture was incubated on ice for 1.5 hours. The following steps were performed at 4 °C or on ice. The culture was transferred to two 50 mL tubes and centrifuged at 5,000 x *g* for 10 minutes. The supernatant was discarded by decanting. Each pellet was resuspended in 50 mL sterile 10% glycerol and centrifuged as above. The supernatant was discarded by decanting. Each pellet was resuspended in 25 mL sterile 10% glycerol and centrifuged at 2,000 x *g* for 10 minutes. The supernatant was discarded by decanting. This wash step was repeated once more. Each pellet was resuspended in 500 µL sterile 10% glycerol and 50 µL aliquots were dispensed into microcentrifuge tubes. These were flash frozen in liquid $N_2$ and stored at -70 °C. Electrocompetent cells were thawed on ice and mixed with 1.0 – 5.0 µL of plasmid DNA. This was transferred to a 2 mm electroporation cuvette and placed on ice for 1 minute. The outside of the cuvette was dried, and it was ensured that no bubbles had formed. The cells were electroporated at 25 µFD, 2.5 KV, 1000 Ω. The cells were mixed with 1 mL MOADC medium and transferred to a 15 mL tube. The cells were incubated at 37 °C for 2 hours with shaking at 250 rpm. 0.1 – 1.0 mL of the culture was plated onto MOADC agar plates (with appropriate antibiotic) and grown for 1-3 days.

**Lydiamycin A bioactivity with *M. smegmatis***

10 mL of MOADC with kanamycin (20 µg/mL) was inoculated with a single colony of *M. smegmatis* $mc^2155$ (containing either empty pJAM2 or pJAM2 harbouring a PDF, Table S4) and grown overnight

at 37 °C with shaking at 250 rpm. The *M. smegmatis* cultures were inoculated into 200 µL of MOADC medium in 96-well microplates to a final concentration of $OD_{600}$ 0.05, in triplicate. All cultures were supplemented with 0.25% TWEEN-80, 0.2% acetamide and kanamycin (20 µg/mL). Each strain was grown either in the presence of methanol (5% v/v) or 50 µM lydiamycin A (in 5% v/v methanol), unless otherwise stated. Cultures were incubated at 37 °C with shaking at 500 rpm. Growth was measured by $OD_{600}$ detection every 30 minutes for 72 hours using a SPECTROstar Nano UV/Vis microplate reader (BMG Labtech). The data were processed in Excel (Microsoft) and Datagraph 5.2 (Visual Data Tools).

#### *In vitro* PDF inhibition assay

PDF from *E. coli* was purified as previously described[27,28]. In short, $His_6$-tagged PDF was expressed in BL21(DE3)pLysS cells (New England Biolabs) and purified by Cobalt-Talon affinity chromatography (Clontech) followed by ion exchange chromatography using a HiTrapQ column (Cytiva). PDF was stored in HEPES buffer (25 mM HEPES, pH 7.5, 70 mM $NH_4Cl$, 30 mM KCl, 7 mM $MgCl_2$, 0.2 mM $CoCl_2$, 1 mM TCEP, and 10% (v/v) glycerol) at -80 °C. To determine the inhibitory effect of lydiamycin on PDF, a colorimetric coupled enzyme assay was carried out using a model peptide, formyl-methionyl-leucyl-p-nitroaniline (fML-pNA, Bachem), as the substrate. PDF (10 nM) was incubated with fML-pNA (20 µM) and *Aeromonas* aminopeptidase (0.8 U/mL) in the presence of lydiamycin A (0-50 µM) in HEPES buffer with 1% DMSO[29,30]. Formation of *p*-nitroaniline upon deformylation was monitored at 405 nm ($\varepsilon_{405}$ = 10,600 $M^{-1}cm^{-1}$) every 5 s at room temperature. The initial velocity was determined by linear regression and the half maximal inhibitory concentration (IC50) was estimated by sigmoidal fitting. The activity of PDF in the presence of 1% DMSO was verified by Michaelis Menten kinetics. The determined parameters of $K_M$ = 27 µM and $k_{cat}$ = 7 $s^{-1}$ are comparable to published values[28,29].

#### Spot-on-lawn assay with *R. fascians*

10 mL of LB was inoculated with a single colony of *R. fascians* and grown overnight at 28 °C with shaking at 250 rpm. This pre-culture was used to inoculate 50 mL of LB (in 250 mL flasks), which was grown at 28 °C with 250 rpm shaking until an $OD_{600}$ of 0.40 was reached. SNA medium was melted and cooled to 42 °C in a water bath. The *R. fascians* culture was added to SNA at a concentration of $OD_{600}$ 0.02 and poured into 120 x 120 mm plates (50 mL each). A concentration series was prepared by serial dilution of pure lydiamycin A in methanol from 600 µg/mL to 1 µg/mL. A positive control of 50 µg/mL kanamycin in water was prepared. 5 µL spots were applied to the plates and allowed to dry. The plates were incubated at 37 °C for 1-3 days and imaged.

#### Root growth assay

A root growth assay was conducted based on methodology in a prior study of *R. fascians*[31]. 120 x 120 mm plates were prepared, each with 50 mL of ½ MS + 3% sucrose + 0.8% agar media. *N. benthamiana* seeds were surface sterilised in 1.5% bleach solution for 30 minutes with strong vortexing. Under aseptic conditions, the seeds were washed in sterile Milli-Q water four times and aliquoted onto sterile filter paper. Twenty seeds were placed on each plate (in a line 2 cm from one edge). Plates were closed with micropore tape and stored in the dark at 4 °C for 3 days to align germination. The plates were placed vertically and grown under 16/8 hour light/dark at 28 °C for 5 days.

10 mL of LB was inoculated with a single colony of *R. fascians* and grown overnight at 30 °C with shaking at 250 rpm. 0.5 mL of culture was centrifuged (4,000 x *g* for 10 minutes), the supernatant

discarded by decanting and the pellet resuspended in 1 mL sterile Milli-Q water. This step was repeated once more, and the pellet resuspended in sterile 100 mM $MgCl_2$ solution to an $OD_{600}$ value of 0.5. The length of the plant root was marked on the plate. Each seedling was inoculated with 4 µL of culture. Each *R. fascians* strain and a mock infection using 4 µL of 100 mM $MgCl_2$ was tested with 60 seedlings. The plates were closed with micropore tape and returned to the same growth conditions for 7 days. The length of each plant root was marked and the distance between the root length at 0 and 7 days-post-inoculation (dpi) was measured. Data were processed using GraphPad Prism.

**LC-MS analysis of seedlings**

Following the root length assay described above, six groups of five seedlings from each condition were transferred separately to 2 mL centrifuge tubes containing 1 mL MeOH and shaken vigorously for 30 minutes. The tubes were centrifuged at 15,871 x *g* for 30 minutes and the resulting supernatants were analysed by LC-MS using instrument settings as described above for mass spectral networking.

**Leafy gall assay**

A leafy gall assay was conducted based on methodology from prior studies of *R. fascians*[32,33]. GA-7 Magenta vessels (Sigma) were prepared, each with 150 mL of ½ MS + 3% sucrose + 0.8% agar media. *N. benthamiana* seeds were surface sterilised in 1.5% bleach solution for 30 minutes with strong vortexing. Under aseptic conditions, the seeds were washed in sterile Milli-Q water four times and aliquoted onto sterile filter paper. One seed was placed into each Magenta vessel, and these were closed and grown with 16/8 hour light/dark at 28 °C for 6 weeks. 10 mL of LB was inoculated with a single colony of *R. fascians* and grown overnight at 30 °C with shaking at 250 rpm. 0.5 mL of culture was centrifuged (4,000 x *g* for 10 minutes), the supernatant discarded by decanting and the pellet resuspended in 1 mL sterile Milli-Q water. This step was repeated once more, and the pellet resuspended in sterile 100 mM $MgCl_2$ solution to an $OD_{600}$ value of 0.5. The meristem of each plant was wounded with sterile forceps and infected with 5 µL of WT *R. fascians* culture, *R. fascians* Δ*lyd* culture or sterile 100 mM $MgCl_2$ solution (n = 9 plants each). The plants were returned to the same growth conditions as above for four weeks and imaged. The galls were also removed, and the masses of gall and plant measured.

**Excised leaf assay**

An excised leaf assay was conducted based on methodology in a prior study of *R. fascians*[14]. GA-7 Magenta vessels (Sigma) were prepared, each with 150 mL of ½ MS + 3% sucrose + 0.8% agar media. *Nicotiana tabacum* cv. Samsun NN seeds were surface sterilised in 1.5% bleach solution for 30 minutes with vortexing. Under aseptic conditions, the seeds were washed in sterile Milli-Q water four times and aliquoted onto sterile filter paper. One seed was placed into each Magenta vessel, and these were closed and grown with 16/8 hour light/dark at 28 °C for 4 weeks. Leaves were aseptically excised and placed onto 120 x 120 mm ½ MS + 3% sucrose + 0.8% agar plates. 10 mL of LB was inoculated with a single colony of *R. fascians* and grown overnight at 30 °C with shaking at 250 rpm. Leaves were infected with 5 µL of WT *R. fascians* culture, *R. fascians* Δ*lyd* culture or sterile 100 mM $MgCl_2$ solution (n = 15 leaves each). The plates were returned to the same growth conditions as above for three weeks and imaged.

**Competition assay**

120 x 120 mm plates were prepared, each with 50 mL of ½ MS + 3% sucrose + 0.8% agar media. *N. benthamiana* seeds were surface sterilised in 1.5% bleach solution for 30 minutes with strong vortexing. Under aseptic conditions, the seeds were washed in sterile Milli-Q water four times and aliquoted onto sterile filter paper. 36 seeds were placed onto each plate. Plates were closed with micropore tape and stored in the dark at 4 °C for 3 days to align germination. The plates were grown under 16/8 hour light at 28 °C for 7 days. 10 mL of LB was inoculated with a single colony of *R. fascians* and grown overnight at 30 °C with shaking at 250 rpm. 0.5 mL of culture was centrifuged (4,000 x *g* for 10 minutes), the supernatant discarded by decanting and the pellet resuspended in 1 mL sterile Milli-Q water. This step was repeated once more, and the pellet resuspended in sterile 100 mM $MgCl_2$ solution to an $OD_{600}$ value of 0.5. 1:1 mixes of *R. fascians* D188-5/WT and *R. fascians* D188-5/Δ*lyd* were prepared. The second competition assay used biological replicates of *R. fascians* WT and Δ*lyd* strains (n = 7 each) from separate colonies, against the same D188-5 biological isolate.

A serial dilution of these inocula was prepared in LB (from $10^{-1}$ to $10^{-8}$ dilution) and 5 µL spots were plated onto both LB and LB + streptomycin plates. Plasmid-free *R. fascians* D188-5 is streptomycin resistant while the competing WT and Δ*lyd* strains are streptomycin sensitive. These plates were incubated at 30 °C for 1-3 days until individual colonies could be counted. The CFU/mL was calculated for each strain. The CFU/mL of the competing strain (WT or Δ*lyd*) was calculated by subtracting the CFU/mL grown under streptomycin selection (count of only D188-5) from the total CFU/mL grown without selection (count of both strains). The same inoculum was used to infect the seedlings. 4 µL of the inoculum (approximately $2x10^5$ cells) was applied to each seedling. In the first experiment, 8 groups of 7 seedlings were infected per condition. In the second experiment, 7 groups of 8 seedlings per timepoint were infected per condition. The plates were grown under 16/8 hour light at 28 °C. In the first experiment, bacteria were isolated at 7 dpi, whereas in the second experiment, bacteria were isolated at 3, 7 and 14 dpi. For both experiments, the seedlings were aseptically transferred to a 2 mL microcentrifuge tube containing 1 mL sterile 100 mM $MgCl_2$ and vigorously mixed for 1 hour. These were serially diluted in LB (from $10^{-1}$ to $10^{-8}$ dilution) and 5 µL spots were plated onto both LB and LB + streptomycin plates. These plates were incubated at 30 °C for 1-3 days until individual colonies could be counted. The CFU/mL were then calculated as described above. The data and statistics were processed in GraphPad Prism, where relative fitness was calculated: ln(test CFU 7dpi/ test CFU 0dpi)/ln(D188-5 CFU 7dpi/D188-5 CFU 0dpi).

**SUPPLEMENTARY TABLES**

**Table S1** Bacterial strains used in this study.

| Strain | Genotype/Description | Application | Reference |
|---|---|---|---|
| *E. coli* DH5α (NEB) | *fhuA2 Δ(argF-lacZ)U169 phoA glnV44 Φ80 Δ(lacZ)M15 gyrA96 recA1 relA1 endA1 thi-1 hsdR17* | Plasmid host for molecular cloning | |
| *M. smegmatis* mc$^2$155 (ATCC 700084) | Wild Type | Mycobacterial reporter strain | 34 |
| *R. fascians* D188 | Wild Type | Lydiamycin producer | 35 |
| *R. fascians* D188-5 | pFiD188-free. Strep$^R$ | Avirulent strain lacking the pFiD188 plasmid. | 35 |
| *R. fascians* D188 Δ*nrp5* (Δ*lyd*) | Disruption in *lydH* | Disrupted lydiamycin production | 14 |
| *M. smegmatis* mc$^2$155::pJAM2-*lydA* | Heterologous expression of *lydA* PDF gene | Testing LydA-mediated lydiamycin resistance | This study |
| *M. smegmatis* mc$^2$155::pJAM2-*MsPDF* | Heterologous expression of *MsPDF* PDF gene | Testing MsPDF-mediated lydiamycin resistance | This study |
| *M. smegmatis* mc$^2$155::pJAM2 | *M. smegmatis* strain harbouring empty pJAM2 plasmid | Control strain for testing lydiamycin activity | This study |

**Table S2** Media used in this study. All percentages are w/v for solid media components.

| Medium | Application | Ingredients |
|---|---|---|
| Lysogeny Broth (LB) | *E. coli* culture, *R. fascians* culture | 1% Bacto-tryptone, 0.5% yeast extract, 1% NaCl, pH 7.0 |
| SOC | *E. coli* transformation recovery medium | 2% tryptone, 0.5% yeast extract, 0.058% NaCl, 0.2% $MgCl_2$, 0.25% $MgSO_4$, 0.36% glucose |
| Middlebrook 7H9 Broth with OADC Enrichment (MOADC liquid) | *M. smegmatis* liquid culture | 0.47% Middlebrook 7H9 broth base, 0.2% glycerol, 10% MOADC growth supplement (after autoclaving) |
| Middlebrook 7H10 Agar with OADC Enrichment (MOADC solid) | *M. smegmatis* solid culture | 1.9% Middlebrook 7H10 agar base, 0.5% glycerol, 10% MOADC growth supplement (after autoclaving) |
| Aurachin production medium (PM)[15] | *R. fascians*, comparative metabolomics | 1% starch, 1% glucose, 1% glycerol. 0.25% corn steep powder, 0.5% bacto peptone, 0.2% yeast extract, 0.1% NaCl, pH 7.3 |
| Rhodostreptomycin PM[16] | *R. fascians*, comparative metabolomics | 1% starch, 2% glucose, 2.5% soytone. 0.4% dry yeast, 0.1% beef extract, 0.005% $K_2HPO_4$, 0.2% NaCl, pH 7.0 |
| Minimal media A (MinA)[17] | *R. fascians*, comparative metabolomics | 1.05% $K_2HPO_4$, 0.45% $KH_2PO_4$, 0.1% $(NH_4)_2SO_4$, 0.05% sodium citrate, 0.025% $MgSO_4$-$7H_2O$, 0.001% thiamine with 20 mM pyruvate and 5 mM histidine |
| Yeast Extract Broth (YEB) | *R. fascians*, comparative metabolomics | 0.5% peptone, 0.1% yeast extract, 0.5% beef extract, 0.5% sucrose, 0.5% $MgSO_4$-$7H_2O$, pH 7.2 |
| Actinonin PM[18] | *R. fascians*, comparative metabolomics | 1% glucose, 1% DIFCO soluble starch, 2% corn liquor step, 2% soy flour, 0.25% $NH_4Cl$, 0.3% NaCl, 0.6% $CaCO_3$, pH 6.2 |
| Matlystatin PM[18] | *R. fascians*, comparative metabolomics | 3% glucose, 7% glycerol, 1% bacto peptone, 1% soy flour, 1% corn steep liquor, 0.5% $MgSO_4$, 0.5% $NH_4NO_3$, 0.5% NaCl |
| Bottromycin PM[19] | *R. fascians*, comparative metabolomics | 1% glucose, 1.5% starch, 0.5% yeast extract, 1% soy flour, 0.5% NaCl, 0.3% CaCO3, pH 7.0 |
| Screening medium 7 (SM7) | *R. fascians*, comparative metabolomics | 2.09% MOPS, 1.5% L-proline, 2% glycerol, 0.25% sucrose, 0.15% L-glutamate, 0.05% NaCl, 0.2% $K_2HPO_4$ with 2 mM $MgSO_4$, 0.2 mM $CaCl_2$, 0.5% trace salts*, pH 6.5 |
| Screening medium 12 (SM12) | *R. fascians*, comparative metabolomics | 1% soy flour, 5% glucose, 0.4% peptone, 0.4% beef extract, 0.1% yeast extract, 0.25% NaCl, 0.5% $CaCO_3$, pH 7.6 |
| Soft Nutrient Agar (SNA) | Spot-on-lawn bioassays | 0.8% Difco Nutrient Broth, 0.7% agar |
| ½ MS + 3% Sucrose | Tobacco plant growth | 0.22% Murashige & Skoog Medium including vitamins, 3% sucrose, 0.8% agar |

* Trace salts (per L in distilled water dissolved in order): 1M $H_2SO_4$ (1 mL), $ZnSO_4.7H_2O$ (0.86 g), $MnSO_4.7H_2O$ (0.223 g), $H_3BO_3$ (62 mg), $CuSO_4.5H_2O$ (0.125 g), $Na_2MoO_4.2H_2O$ (48 mg) $CoCl_2.6H_2O$ (48 mg) $FeSO_4.7H_2O$ (1.8 g), KI (83 mg)

**Table S3** Primers used in this study.

| Primer Name | Sequence (5' to 3') | Application | Restriction Site |
|---|---|---|---|
| pJAM LydA For | GATACAGGATCCATGCCTGTCTCTGAA CTTCTGC | Amplification of *lydA* for pJAM2-*lydA* construction | BamHI |
| pJAM LydA Rev | GATACATCTAGACTATCGTGTTGGCCA ATCGTTG | | XbaI |
| pJAM MsPDF For | CATGCCCGAGGTAGTTTTCGGATCCAT GGCCGTCGTCCCGATCC | Amplification of *M. smegmatis* housekeeping PDF for pJAM2-*MsPDF* construction | Gibson |
| pJAM MsPDF Rev | AGTGGTGGTGGTGGTGGTGTCTAGATC AGTGCCCGAACGGATCG | | Gibson |

**Table S4** Vectors used in this study.

| Vector | Features | Application | Resistance Marker |
|---|---|---|---|
| pJAM2 [26] | Acetamide-inducible promotor. C-terminal His tagging. | Induced expression in *M. smegmatis* | Kanamycin |
| pJAM2-*lydA* | pJAM2 with *lydA* insert. C-terminal His-tag. | Expression of lydiamycin BGC PDF in *M. smegmatis* | Kanamycin |
| pJAM2-*MsPDF* | pJAM2 with *MsPDF* (WP_011727226.1) insert. C-terminal His-tag. | Expression of *M. smegmatis* PDF in *M. smegmatis* | Kanamycin |

**Table S5** Accessions of mutases from EFI-EST analysis associated with the chemophore clade.

| Representative mutase accession | Mutase accessions grouped using EFI-EST[a] |
|---|---|
| NNN35785.1<br>*Streptomyces* sp. S3 | |
| WP_104814753.1<br>*Kitasatospora* sp. MMS16-BH015 | |
| KY906183.1<br>*Streptomyces* sp. ATCC 14903 | WP_086815435.1<br>*Streptomyces cacaoi* NPDC 016406 |
| | WP_159787793.1<br>*Streptomyces* sp. NHF165 |
| | QNF54062.1<br>*Streptomyces* sp. KY2 |
| | QHF95772.1<br>*Streptomyces* sp. NHF165 |
| WP_210837750.1<br>*Micromonospora* sp. C31 | |
| WP_019634562.1<br>*Actinomadura atramentaria* | |
| ALC26898.1<br>*Streptomyces* sp. CFMR 7 | MBB4761836.1<br>*Actinoplanes digitatis* DSM 43149 |
| | ALC26898.1<br>*Streptomyces* sp. CFMR 7 |
| | MBA3364174.1<br>MAG[b]: Actinomycetota bacterium |
| | WP_176725504.1<br>*Streptomyces* sp. DvalAA-19 |
| | WP_189742219.1<br>*Streptomyces aureoverticillatus* |
| | WP_223771548.1<br>*Streptomyces huiliensis* |
| AET25252.1<br>*Rhodococcus fascians* D188[c] | |
| AMY56278.1<br>*Rhodococcus fascians* D188[c] | OZE97895.1<br>*Rhodococcus* sp. 15-2388-1-1a |
| | OZE22244.1<br>*Rhodococcus* sp. 05-2255-1e |
| | OZF08685.1<br>*Rhodococcus* sp. 14-2686-1-2 |
| | OZF10685.1<br>*Rhodococcus fascians* 14-2632-D2 |
| | OZC51310.1<br>*Rhodococcus* sp. 06-621-2 |
| | OZC60729.1<br>*Rhodococcus* sp. 06-469-3-2 |
| | OZC94541.1<br>*Rhodococcus* sp. 06-412-2C |
| | OZD10640.1<br>*Rhodococcus* sp. 06-221-2 |
| | OZD31149.1<br>*Rhodococcus* sp. 06-156-3 |
| | OZD86423.1<br>*Rhodococcus* sp. 05-2256-B3 |
| | OZE41003.1<br>*Rhodococcus* sp. 05-2254-4 |
| | OZE57960.1<br>*Rhodococcus* sp. 02-925g |
| | OZE73377.1<br>*Rhodococcus* sp. 15-649-2-2 |

| | |
|---|---|
| | OZE75211.1<br>*Rhodococcus* sp. 15-649-1-2 |
| WP_063760750.1<br>*Streptomyces aureocirculatus* | |
| MYU63353.1[d]<br>*Streptomyces* sp. SID69 | |
| MYY82164.1<br>*Streptomyces* sp. SID335 | MYZ12566.1<br>*Streptomyces* sp. SID337 |
| | NDZ85767.1<br>*Streptomyces* sp. SID10115 |
| | NEA03361.1<br>*Streptomyces* sp.. SID10116 |
| | NEB46693.1<br>*Streptomyces* sp. SID339 |
| QES31823.1<br>*Streptomyces venezuelae* ATCC 14583 | MBA2638190.1<br>MAG[b]: Solirubrobacterales bacterium |

a. NCBI accessions of the proteins from the EFI-EST analysis that group with the representative accession. See Materials and Methods for details of settings.
b. MAG = metagenome-assembled genome.
c. Two alternative annotations for the D188 mutase. The separate grouping by EFI-EST likely relates to one accession being annotated with a different start codon.
d. Partial sequence of only 71 AA.

**Table S6** Predicted functions of the proteins encoded in the *R. fascians* D188 *lyd* BGC.

| **Gene name** | **Protein accession number** | **Pfam domain** | **Predicted function** | **Size (AA)** | **Size (kDa)** |
|---|---|---|---|---|---|
| *lydA* | WP_015586162.1 | PF01327 | Peptide deformylase | 188 | 20.71 |
| *lydB* | AET25245.1 | PF00196 | LuxR-type regulator | 226 | 24.86 |
| *lydC* | WP_015586164.1 | PF01040 | UbiA family prenyltransferase | 282 | 29.44 |
| *lydD* | WP_015586165.1 | PF03621 | MbtH-like | 103 | 11.50 |
| *lydE* | WP_015586166.1 | PF13434 | KtzI-like | 424 | 46.95 |
| *lydF* | WP_202902192.1 | PF04299 | KtzT-like | 218 | 24.51 |
| *lydG* | WP_015586168.1 | PF01494 | Monooxygenase | 394 | 42.68 |
| *lydH* | WP_015586169.1 | Multiple | NRPS | 4,925 | 537.17 |
| *lydI* | AET25252.1 | PF01642 | Hexylmalonyl-CoA mutase (α subunit) | 461 | 49.76 |
| *lydJ* | AET25253.1 | PF00550 | NRPS | 99 | 11.06 |
| *lydK* | WP_015586172.1 | PF00501 | NRPS | 527 | 56.99 |
| *lydL* | WP_015586173.1 | PF00668 | NRPS | 614 | 67.49 |
| *lydM* | WP_015586174.1 | PF08240 | Crotonyl-CoA carboxylase/reductase (CCR) | 440 | 47.68 |
| *lydN* | WP_015586175.1 | PF02310 | Hexylmalonyl-CoA mutase (β subunit) | 151 | 16.37 |
| *lydO* | WP_015586176.1 | PF03308 | MeaB-like | 251 | 26.66 |
| *lydP* | WP_015586177.1 | PF13669 | Hexylmalonyl-CoA epimerase | 144 | 15.89 |

**Table S7** Lydiamycin A NMR assignment (carbon and proton) and observed 2D cross-peaks (HMBC, COSY, 1,1-ADEQUATE). All NMR recorded in $CDCl_3$. Atom numbering of the revised structure is used (Figure S10).

| Position | $\delta_C$ ppm | $\delta_H$ ppm (multiplicity, *J* in Hz) | HMBC (C to H) | COSY | 1,1-ADEQUATE |
|---|---|---|---|---|---|
| **1** | 169.8 | | 16, 17 | | 2 |
| **2** | 52.7 | 5.32, m | 3, 4 | 3 | 3 |
| **3** | 24.4 | 2.38, 1.79, m | 4, 5, 2 | | |
| **4** | 21.5 | 1.63, 1.59, m | 2, 5, 5-NH | 3, 5 | 5 |
| **5** | 47.1 | 3.16, br d (13.4)<br>2.78, qd (13.1, 3.5) | 3, 4 | 4, 5-NH | |
| **5-NH** | | 4.33, br d | | | |
| **6** | 174.4 | | 2, 5-NH, 7, 7-NH, 8 | | 7 |
| **7** | 50.5 | 4.86, qd (10.4, 7.3) | 7-NH, 8 | 7-NH, 8 | 8 |
| **7-NH** | | 7.31, d (10.4) | | 7 | |
| **8** | 18.3 | 1.46, d (7.3) | 7 | 7, 7-NH | 7 |
| **9** | 169.0 | | 7, 7-NH, 10, 11 | | 10 |
| **10** | 51.4 | 4.35, m | 7-NH, 10-NH, 11, 12 | 10-NH, 11 | 11 |
| **10-NH** | | 7.47, d (8.9) | | 10 | |
| **11** | 36.5 | 1.77, m | 10, 10-NH, 12 | 10 | 10 |
| **12** | 24.6 | 1.63, m | 10 | | 11, 13, 14 |
| **13** | 22.8 | 0.94, d (6.6) | 11, 12 | 12 | |
| **14** | 22.3 | 0.89,(6.6) | 12 | | |
| **15** | 169.4 | | 10, 10-NH, 16, 16-NH, 17 | | 16 |
| **16** | 50.7 | 5.31, m | 16-NH, 17 | 16-NH, 17 | 17 |
| **16-NH** | | 8.31, d (9.9) | | 16 | |
| **17** | 68.3 | 4.79, dd (11.4, 4.3)<br>4.06, d (11.4) | 16 | 16 | 16 |
| **18** | 171.0 | | 16, 16-NH, 19, 20 | | 19 |
| **19** | 55.6 | 4.68, t (5.0) | 20, 21 | 20, 21 | 20 |
| **20** | 19.4 | 2.38, 1.95, m | 19, 21, 22 | 19, 21, 22 | 19 |
| **21** | 20.7 | 2.17, 1.95, m | 19, 20, 22 | 19, 20, 22 | 22 |
| **22** | 142.7 | 6.93, m | 20, 21 | 20, 21 | 21 |
| **23** | 177.4 | | 19, 24, 25, 30 | | 24 |
| **24** | 46.0 | 3.05, m | 25, 26, 30 | 25, 30 | 25, 30 |
| **25** | 31.7 | 1.75, 1.54, m | 24, 30 | | 24 |
| **26** | 26.9 | 1.37, 1.26, m | 24, 25, 27 | 25 | 25 |
| **27** | 31.5 | 1.29, 1.26, m | 25, 26 | 25, 26, 29 | 28 |
| **28** | 22.4 | 1.28, m | 26 | 26, 29 | 29 |
| **29** | 14.0 | 0.87, t (6.9) | 27, 28 | | 28 |
| **30** | 36.6 | 3.33, dd (17.0, 12.3)<br>2.63, dd (17.0, 5.5) | 24 | 24 | 24 |
| **31** | 176.0 | | 30 | | 30 |
| **31-OH** | | 11.41, br s | | | |

**Table S8** Comparison between published lydiamycin A $^1H$ NMR spectra and data collected in this current study. All NMR recorded in $CDCl_3$.

| **Position in original structure (from ref. 36)** | **Position in revised structure[a] (see Fig. S10)** | **Reported ref. 36 $\delta_H$ (*J* in Hz)** | **Reported ref. 37 $\delta_H$ (*J* in Hz)** | **Observed $\delta_H$ (*J* in Hz)** |
|---|---|---|---|---|
| **1** | **1** | | | |
| **2** | **2** | 5.30, m | 5.28, m | 5.32, m |
| **3** | **3** | 2.38, 1.77, m | 2.35, 1.77, m | 2.38, 1.79, m |
| **4** | **4** | 1.63, 1.59, m | 1.60, m | 1.63, 1.59, m |
| **5** | **5** | 3.14, m, 2.77, qd (12.0, 3.0) | 3.12, br d (13.5), 2.75, br dq (13.5, 3.8) | 3.16, br d (13.4) 2.78, qd (13.1, 3.5) |
| **5-NH** | **5-NH** | 4.32, d (12.0) | 4.31, br d | 4.33, br d |
| **6** | **6** | | | |
| **7** | **7** | 4.85, qd (10.5, 7.0) | 4.83, qd (10.5, 7.25) | 4.86, qd (10.4, 7.3) |
| **7-NH** | **7-NH** | 7.30, d (10.5) | 7.27. d (10.5) | 7.31, d (10.4) |
| **8** | **8** | 1.44, d (7.0) | 1.43, d (7.25) | 1.46, d (7.3) |
| **9** | **9** | | | |
| **10** | **10** | 4.35, td (8.5, 8.5) | 4.33, m | 4.35, m |
| **10-NH** | **10-NH** | 7.46, d (8.5) | 7.43, d (7.8) | 7.47, d (8.9) |
| **11** | **11** | 1.78, m | 1.77, m | 1.77, m |
| **12** | **12** | 1.65, m | 1.63, m | 1.63, m |
| **13** | **13** | 0.93, d (6.5) | 0.93, d (6.5) | 0.94, d (6.6) |
| **14** | **14** | 0.88, d (6.5) | 0.87, d (7.8) | 0.89,(6.6) |
| **15** | **15** | | | |
| **16** | **16** | 5.31, m | 5.28, m | 5.31, m |
| **16-NH** | **16-NH** | 8.30, d (10.0) | 8.25, d (9.85) | 8.31, d (9.9) |
| **17** | **17** | 4.77, dd (11.5, 4.5), 4.04, d (11.5) | 4.75, dd (11.4, 4.4), 4.03, br d (11.4) | 4.79, dd (11.4, 4.3) 4.06, d (11.4) |
| **18** | **18** | | | |
| **19** | **19** | 4.67, t (5.0) | 4.65, br t (5.7) | 4.68, t (5.0) |
| **20** | **20** | 2.35, 1.92, m | 2.35, 1.93, m | 2.38, 1.95, m |
| **21** | **21** | 2.17, 1.95 m | 2.15, 1.92, m | 2.17, 1.95, m |
| **22** | **22** | 6.92, m | 6.90, br s | 6.93, m |
| **23** | **31** | | | |
| **24** | **30** | 3.32, dd (17.0, 12.0), 2.62, dd (17.0, 5.5) | 3.30, dd (16.8, 12.2), 2.60, dd (16.8, 5.3) | 3.33, dd (17.0, 12.3) 2.63, dd (17.0, 5.5) |
| **25** | **24** | 3.04, m | 3.02, m | 3.05, m |
| **26** | **25** | 1.72, 1.53, m | 1.71, 1.52, m | 1.75, 1.54, m |
| **27** | **26** | 1.35, 1.25, m | 1.35, 1.24, m | 1.37, 1.26, m |
| **28** | **27** | 1.29, 1.25, m | 1.27-1.26, m | 1.29, 1.26, m |
| **29** | **28** | 1.27, m | 1.27-1.26, m | 1.28, m |
| **30** | **29** | 0.86, t (7.0) | 0.85, t (6.5) | 0.87, t (6.9) |
| **31** | **23** | | | |
| **31-OH** | **31-OH** | 11.37, br s | | 11.40, br s |

[a]Atom numbering that differs with the prior assignment is highlighted in red.

**Table S9** Comparison between published lydiamycin A $^{13}C$ NMR spectra and data collected in this current study. All NMR recorded in $CDCl_3$.

| Position in original structure (from ref. [36]) | Position in revised structure[a] (see Fig. S10) | Reported ref. [36] δc, multi. | Reported ref. [37] δc, multi. | Observed δc |
|---|---|---|---|---|
| 1 | 1 | 169.8, qC | 169.1, qC | 169.8 |
| 2 | 2 | 52.7, CH | 52.7, CH | 52.7 |
| 3 | 3 | 24.4, CH2 | 24.4, CH2 | 24.4 |
| 4 | 4 | 21.4, CH2 | 21.4, CH2 | 21.5 |
| 5 | 5 | 47.1 CH2 | 47.1, CH2 | 47.1 |
| 6 | 6 | 174.3, qC | 174.3, qC | 174.4 |
| 7 | 7 | 50.5, CH | 50.5, CH | 50.5 |
| 8 | 8 | 18.3 | 18.2 | 18.3 |
| 9 | 9 | 169.0, qC | 169.3, qC | 169.0 |
| 10 | 10 | 51.4, CH | 51.4, CH | 51.4 |
| 11 | 11 | 36.4, CH2 | 36.6, CH2 | 36.5 |
| 12 | 12 | 24.6, CH | 24.6, CH | 24.6 |
| 13 | 13 | 22.8, CH3 | 22.7, CH3 | 22.8 |
| 14 | 14 | 22.3, CH3 | 22.2, CH3 | 22.3 |
| 15 | 15 | 169.3, qC | 169.7, qC | 169.4 |
| 16 | 16 | 50.6, CH | 50.7, CH | 50.7 |
| 17 | 17 | 68.3, CH2 | 68.2, CH2 | 68.3 |
| 18 | 18 | 171.0, qC | 170.9, qC | 171.0 |
| 19 | 19 | 55.6, CH | 55.6, CH | 55.6 |
| 20 | 20 | 19.4, CH2 | 19.3, CH2 | 19.4 |
| 21 | 21 | 20.6, CH2 | 20.6, CH2 | 20.7 |
| 22 | 22 | 142.7, CH | 142.7, CH | 142.7 |
| 23 | 31 | 175.9, qC | 175.9, qC | 176.0 |
| 24 | 30 | 36.6, CH2 | 36.5, CH2 | 36.6 |
| 25 | 24 | 46.0, CH | 46.0, CH | 46.0 |
| 26 | 25 | 31.6, CH2 | 31.6, CH2 | 31.7 |
| 27 | 26 | 26.9, CH2 | 26.8, CH2 | 26.9 |
| 28 | 27 | 31.5, CH2 | 31.5, CH2 | 31.5 |
| 29 | 28 | 22.4, CH2 | 22.4, CH2 | 22.4 |
| 30 | 29 | 14.0, CH3 | 13.9, CH3 | 14.0 |
| 31 | 23 | 177.3, qC | 177.3 qC | 177.4 |

[a]Atom numbering that differs with the prior assignment is highlighted in red.

### SUPPLEMENTARY FIGURES

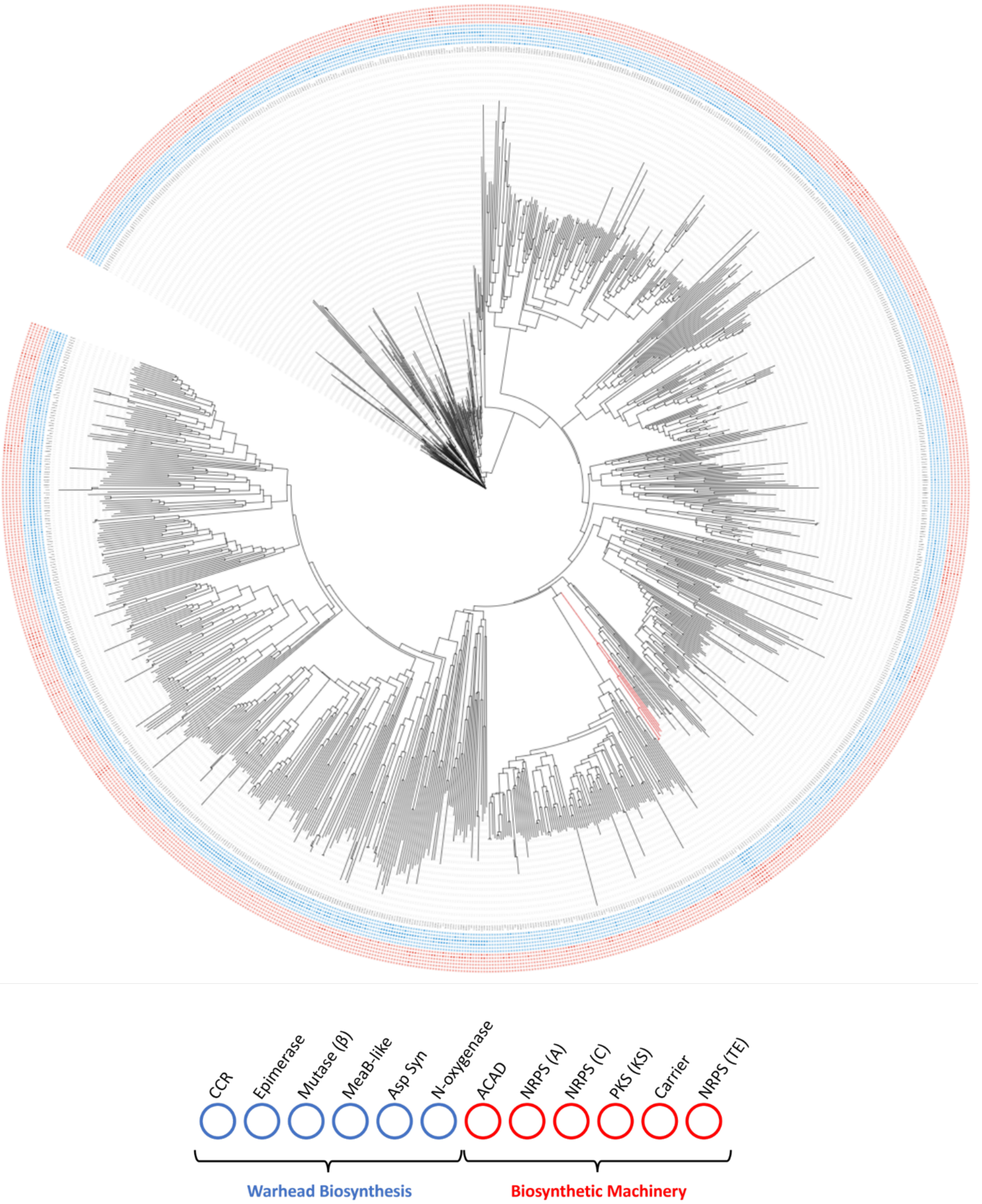


**Figure S1** Phylogenetic tree of actinobacterial methylmalonyl-CoA mutases (α subunit). The tree is annotated with the co-association analysis of pfam domains that are proposed to be involved in HPS analysis or assembly line biosynthesis (NRPS or PKS), and are encoded within 15 genes each side of the mutase gene. A filled blue circle indicates the presence of a proposed HPS chemophore gene. A filled red circle indicates the presence of an NRPS or PKS gene. The 'chemophore clade' is coloured in red. Abbreviations: CCR = crotonyl-CoA carboxylase/reductase, asparagine synthetase, NRPS = non-ribosomal peptide synthetase (adenylation, condensation), PKS = polyketide synthase (ketosynthase), ACAD = acyl-CoA dehydrogenase (proposed to be involved in the dehydrogenation of the carboxy terminus of matlystatin), TE= thioesterase.

A

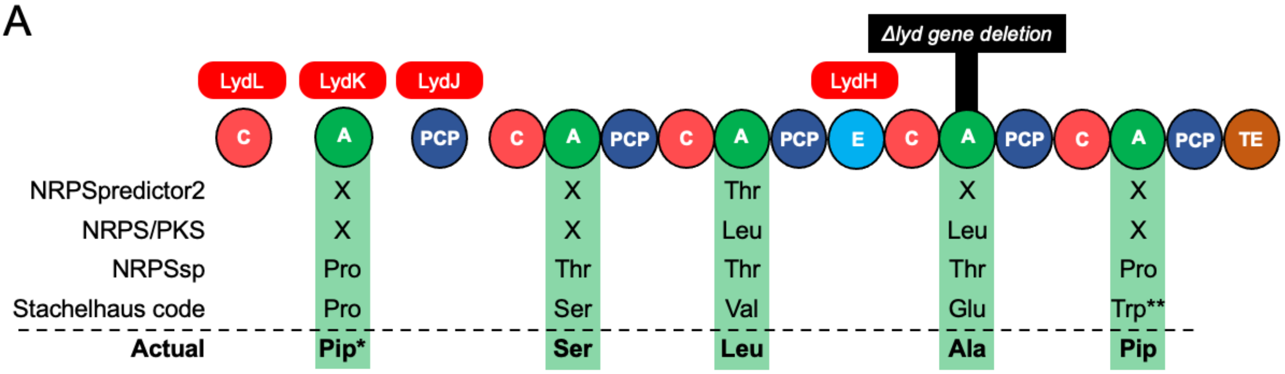


B

| Adenylation Domain | Stachelhaus sequence | | | | | | | | | | Nearest Stachelhaus code (% match) | Known Substrate | |
|---|---|---|---|---|---|---|---|---|---|---|---|---|---|
| LydH M1 | D | V | W | S | V | A | M | V | H | K | Ser (90%) | Ser | |
| LydH M2 | D | F | W | T | L | G | A | V | Y | K | Val (70%) | Leu | |
| LydH M3 | D | V | W | T | F | G | S | V | Y | K | Glu (70%) | Ala | |
| LydH M4 | D | V | W | N | F | G | G | F | A | K | N-(1,1-dimethyl-1-allyl)Trp (70%) | Pip | |
| LydK | D | V | Q | F | N | A | H | M | V | K | Pro (80%) | Pip* | ▲ |
| MatJ | D | V | Q | F | N | A | H | M | V | K | Pro (80%) | Pip | ▲ |
| MatO | D | A | W | F | L | G | N | I | I | K | Leu (80%) | Leu | |
| ActH | D | F | W | N | I | G | M | V | F | K | Thr (90%) | Val | |
| ActE | D | V | Q | F | V | A | H | V | V | K | Pro (90%) | Pro | |

**Figure S2** The domain specificity of the *R. fascians lyd* NRPS. A. Schematic of NRPS and predicted domain architecture. The predicted adenylation specificity using four different tools is denoted[9–12]. The domain that is disrupted in the Δ*lyd* mutant is highlighted. Abbreviations: A = adenylation, C = condensation, PCP = peptide carrier protein, E = epimerase, TE = thioesterase, Pip = piperazic acid. *Lydiamycin A features a dehydropiperazic acid predicted to be incorporated by LydK but the timing of dehydrogenation is not known. **See panel B for the full Trp-like molecule. B. Stachelhaus specificity sequences of *lyd* adenylation modules identified by antiSMASH 6.2[9]. Adenylation domains from the related actinonin and matlystatin BGCs are included as references. Amino acids are colour-coded using the RasMol Shapely scheme and identical specificity codes are denoted by triangles. Abbreviations: Pip = piperazic acid, M = module.

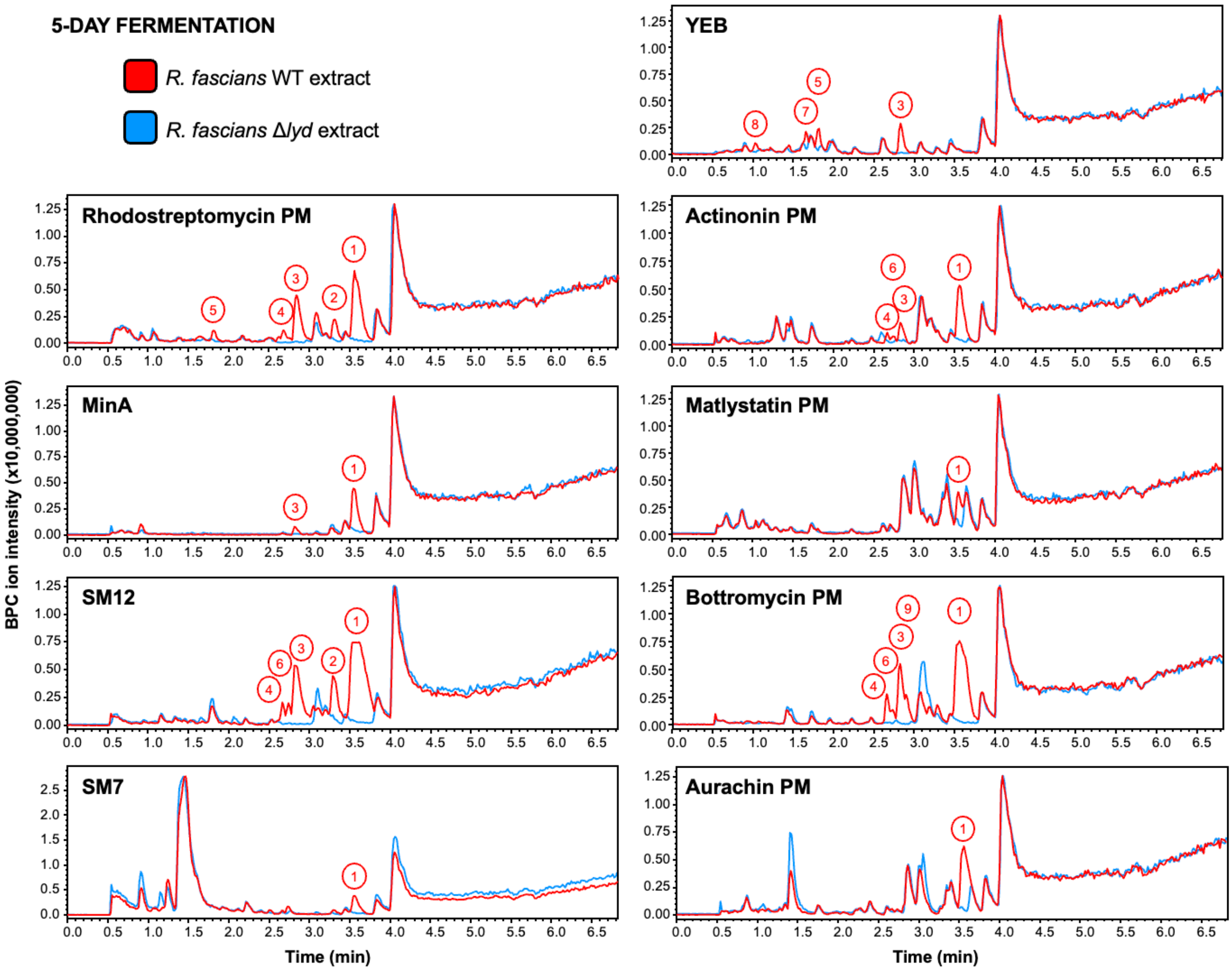


**Figure S3** LC-MS chromatograms comparing *R. fascians* WT and Δ*lyd* strains fermented in different media for 5 days. BPC = base peak chromatogram; PM = production medium. Numbers highlight metabolites seen in WT extracts but not in Δ*lyd* and are listed in Figure S5.

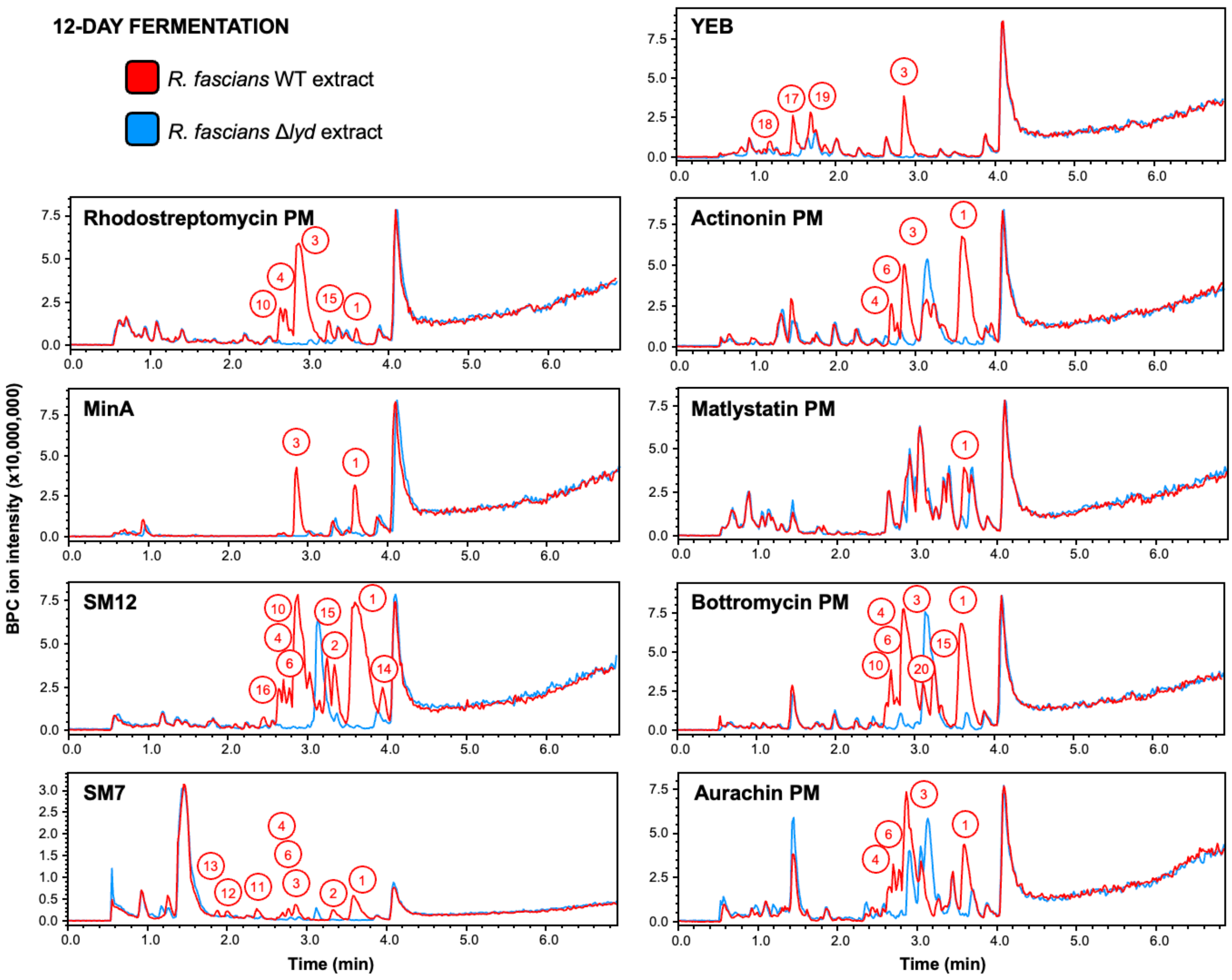


**Figure S4** LC-MS chromatograms comparing *R. fascians* WT and Δ*lyd* strains fermented in different media for 12 days. BPC = base peak chromatogram; PM = production medium. Numbers highlight metabolites seen in WT extracts but not in Δ*lyd* and are listed in Figure S5.

A

| Compound number | Retention time / min | *m/z* | Compound number | Retention time / min | *m/z* |
|---|---|---|---|---|---|
| 1 | 3.55 | 664.37 | 11 | 2.37 | 580.31 |
| 2 | 3.31 | 650.35 | 12 | 2.00 | 577.27 |
| 3 | 2.84 | 682.38 | 13 | 1.88 | 605.26 |
| 4 | 2.67 | 570.30 | 14 | 3.94 | 700.36 |
| 5 | 1.81 | 530.30 | 15 | 3.25 | 664.37 |
| 6 | 2.74 | 689.35 | 16 | 2.44 | 654.35 |
| 7 | 1.66 | 659.34 | 17 | 1.48 | 553.34 |
| 8 | 1.04 | 603.31 | 18 | 1.16 | 594.33 |
| 9 | 2.90 | 703.36 | 19 | 1.70 | 659.34 |
| 10 | 2.64 | 668.36 | 20 | 3.10 | 655.28 |

B

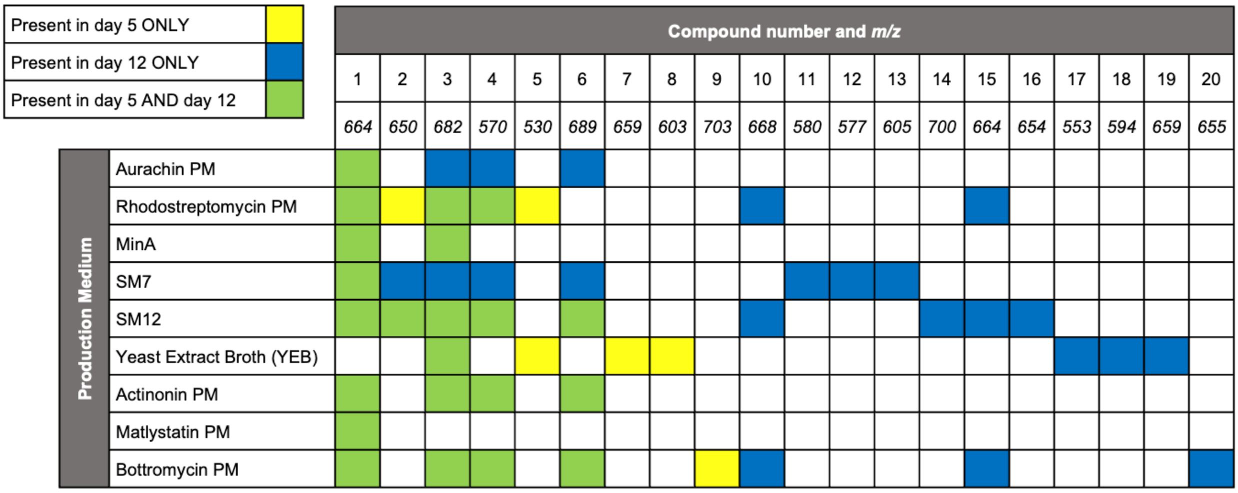


**Figure S5** Summary of potential metabolites associated with the *lyd* BGC in different production media. A. List of metabolites observed in Figures S3 and S4. B. Matrix of metabolites versus production medium at different fermentation times.

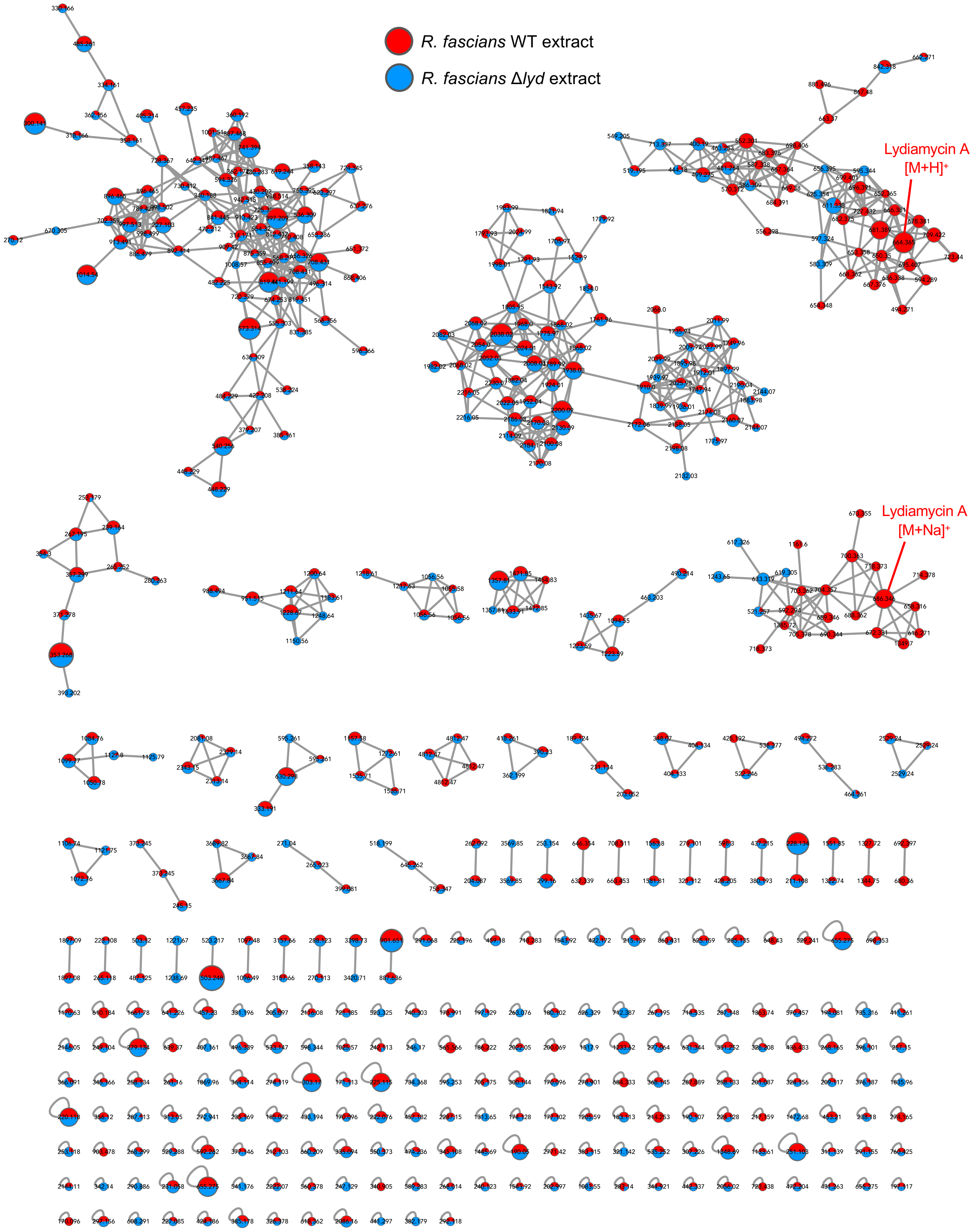


**Figure S6** Mass spectral networking of extracts of *R. fascians* WT and Δ*lyd* strains from 5-day SM12 and YEB cultures. Molecular networking was performed with GNPS[21] and visualised using Cytoscape[4]. The node pie charts reflect the relative number of MS/MS spectra for each molecule between *R. fascians* WT and Δ*lyd* samples. Nodes are annotated with parent masses. The diameter of each node is relative to the total number of MS/MS spectra for each molecule: minimum (2 spectra) = 25 node size; maximum (76 spectra) = 75 node size.

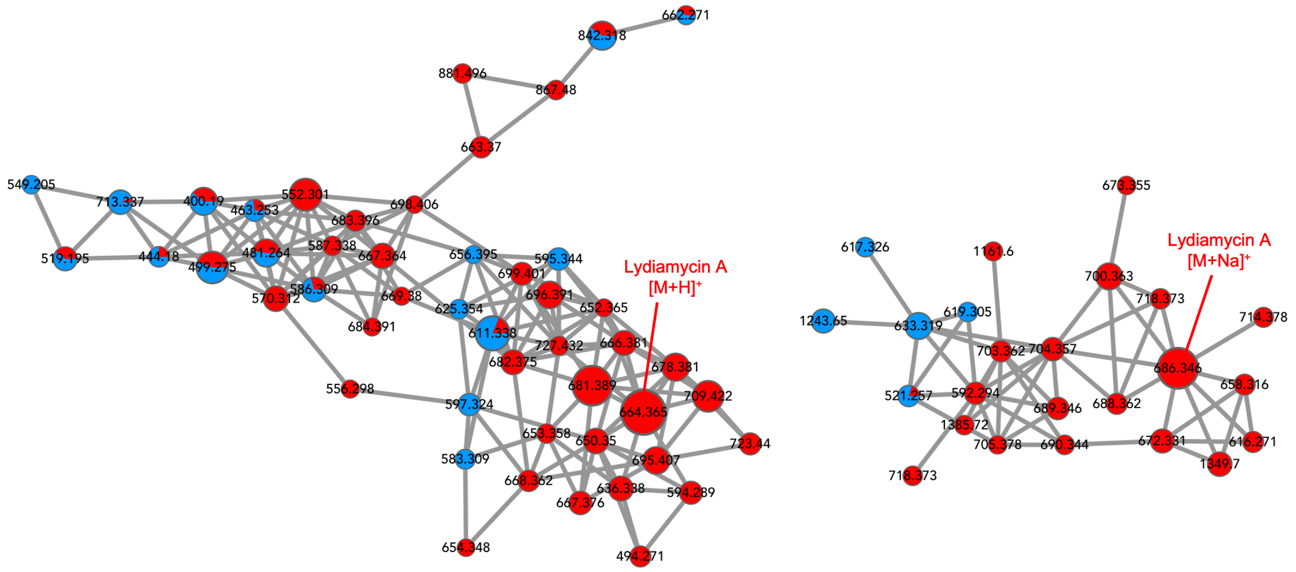


**Figure S7** Magnified detail of networks 3 and 4 from Figure S6. The nodes that feature molecules present in Δ*lyd* may reflect low amounts of alternative products of the pathway arising from the in-frame deletion in the *lydH* gene corresponding to the NRPS adenylation domain (see Figure S2).

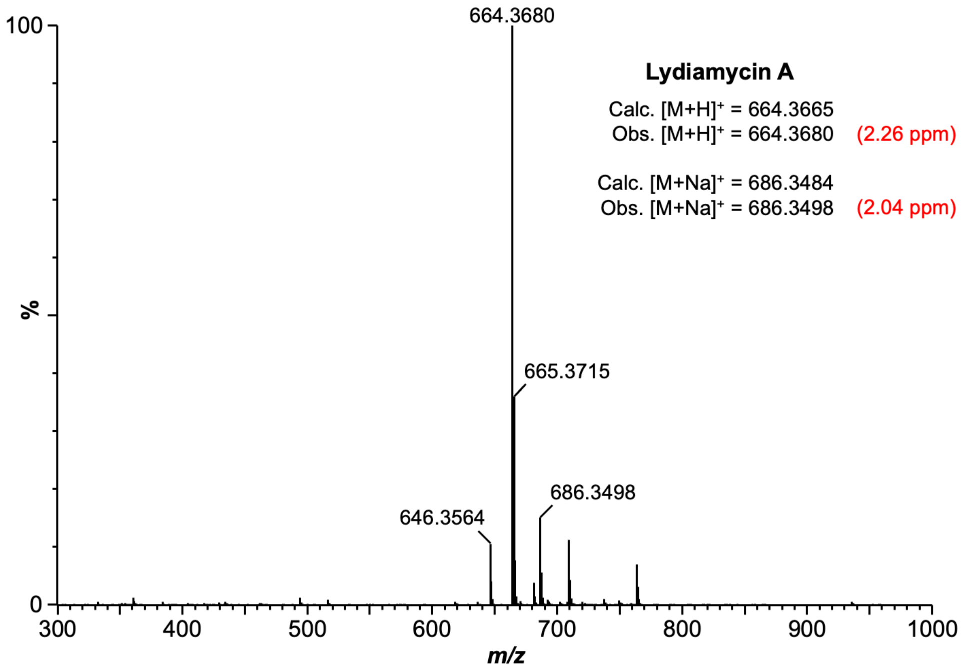


**Figure S8** HR-MS spectrum of lydiamycin A, the major *lyd* BGC product. Data acquired on a Synapt G2-Si mass spectrometer.

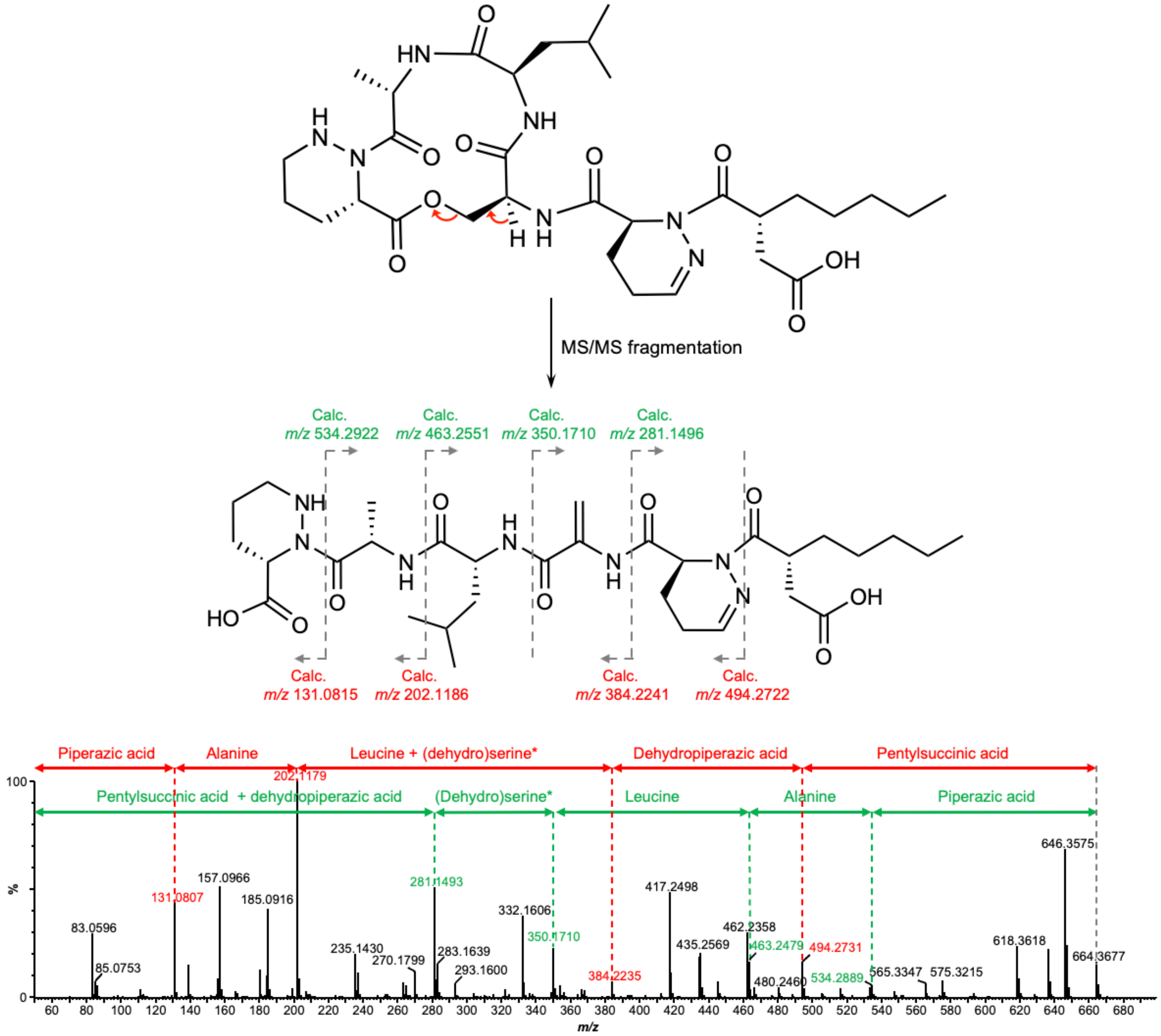


**Figure S9** MS/MS fragmentation of lydiamycin A acquired on a Synapt G2-Si mass spectrometer. Observed *y*-ion and *b*-ion masses are annotated at each amide bond. Asterisk denotes that the serine residue is predicted to be dehydrated during molecule linearisation under MS/MS conditions.

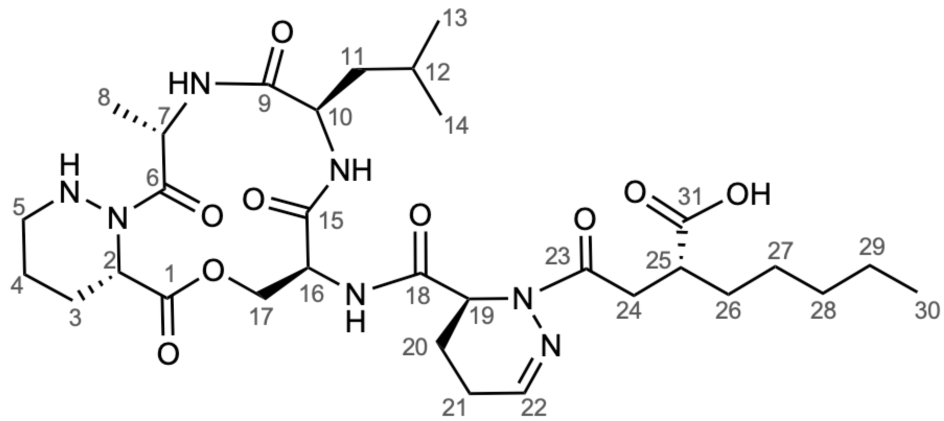


**Lydiamycin A structure and numbering from Hwang *et al*.**

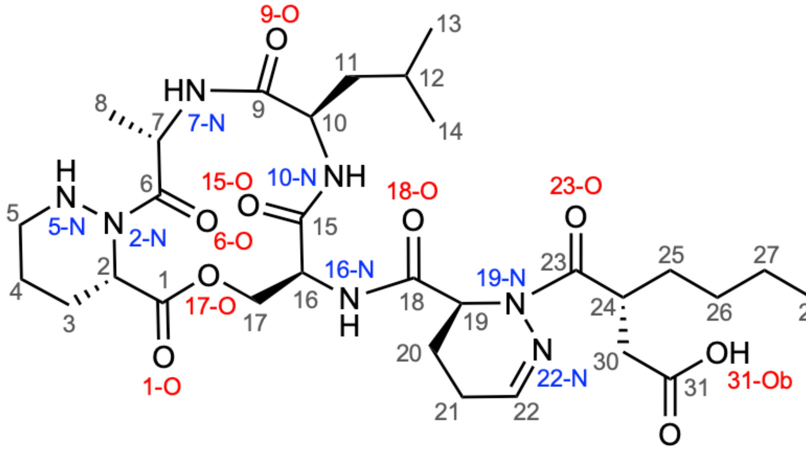


**Revised lydiamycin A structure and numbering (this work)**

**Figure S10** Structures and atom numbering for lydiamycin A from the study of Hwang *et al*.[36] (top) and the revised version from this current work (bottom).

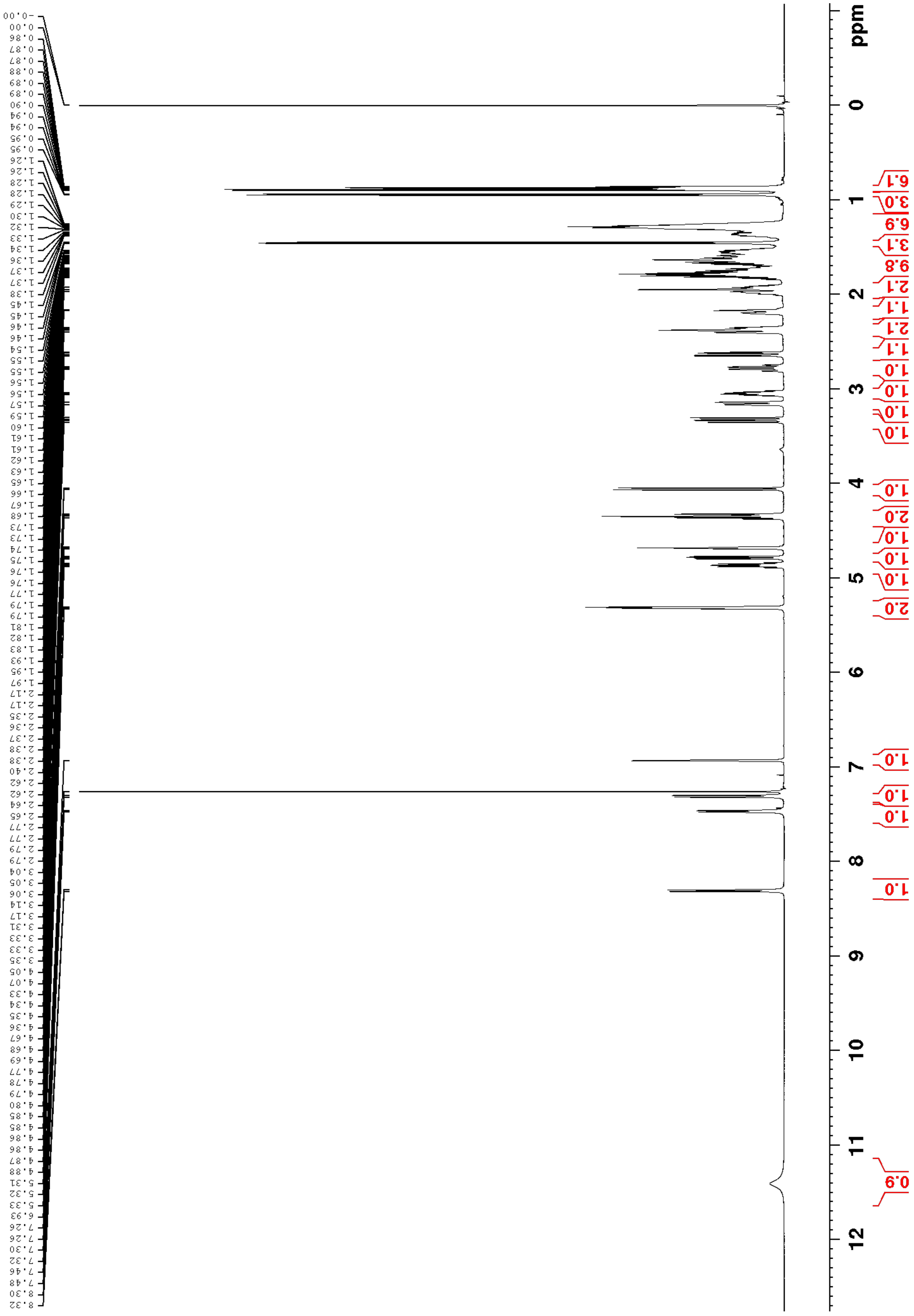

**Figure S11** Lydiamycin A $^{1}$H NMR spectrum (600 MHz, $CDCl_3$, 298 K).

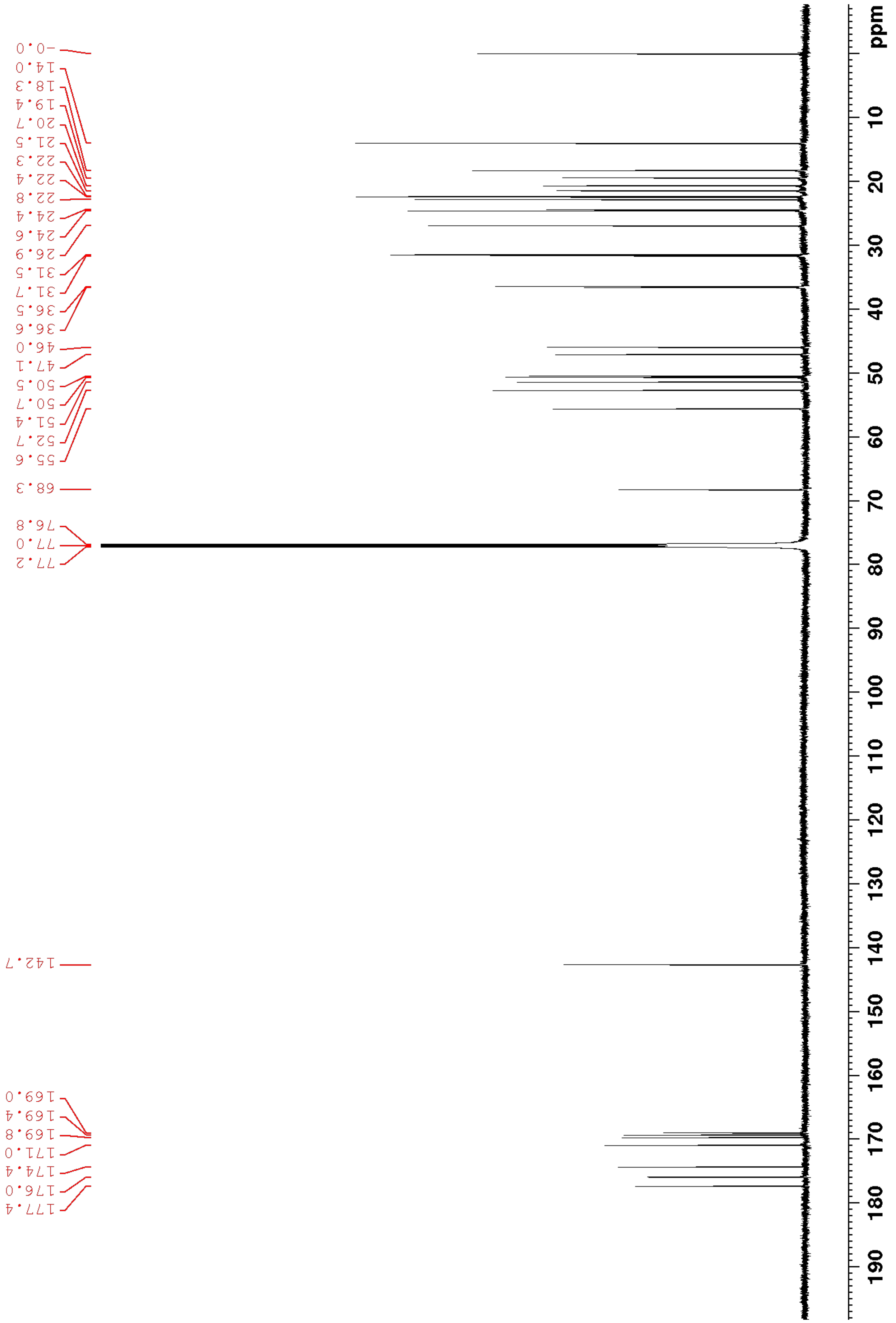


**Figure S12** Lydiamycin A $^{13}C$ NMR spectrum (150 MHz, $CDCl_3$, 298 K).

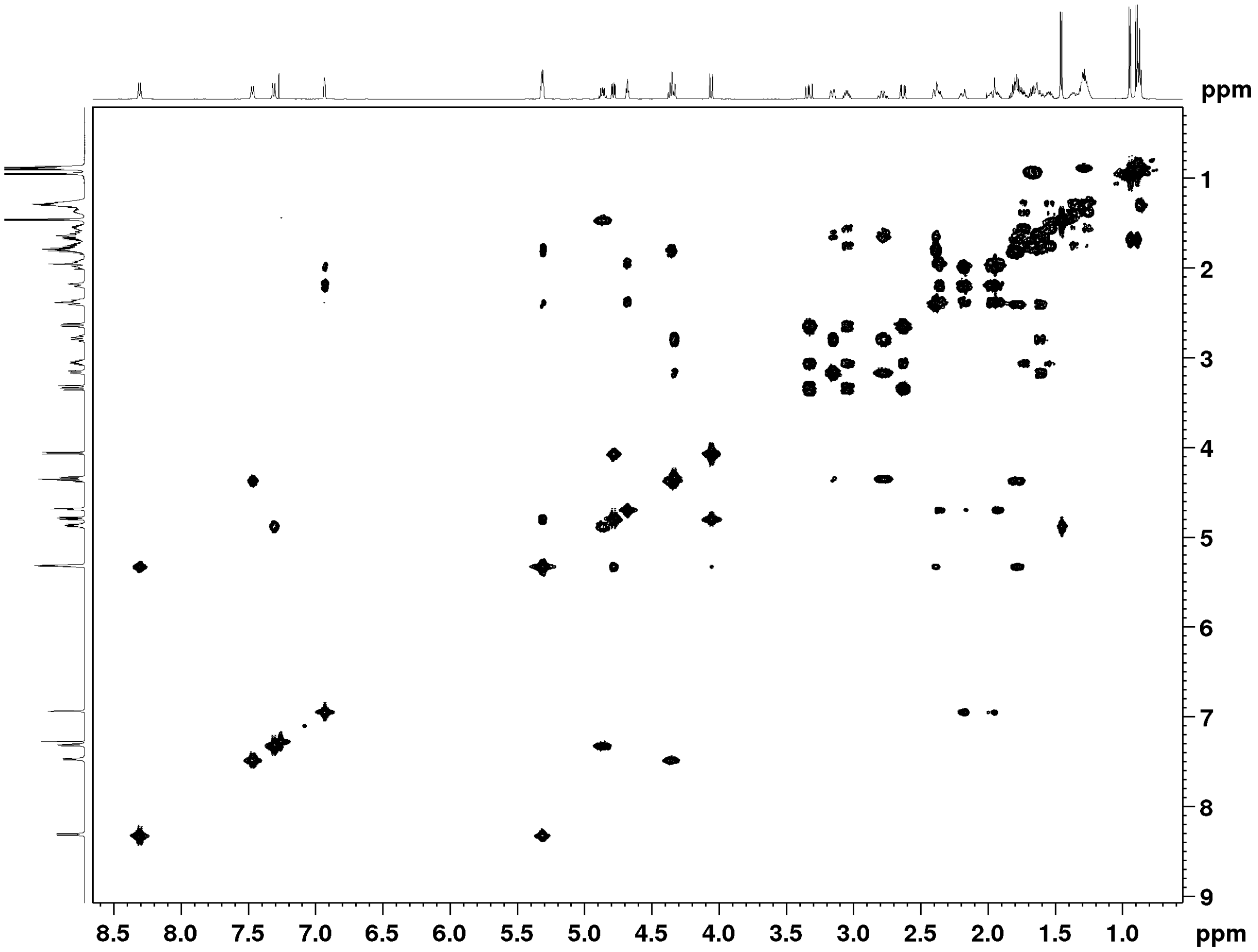


**Figure S13** Lydiamycin A COSY NMR spectrum ($CDCl_3$, 298 K).

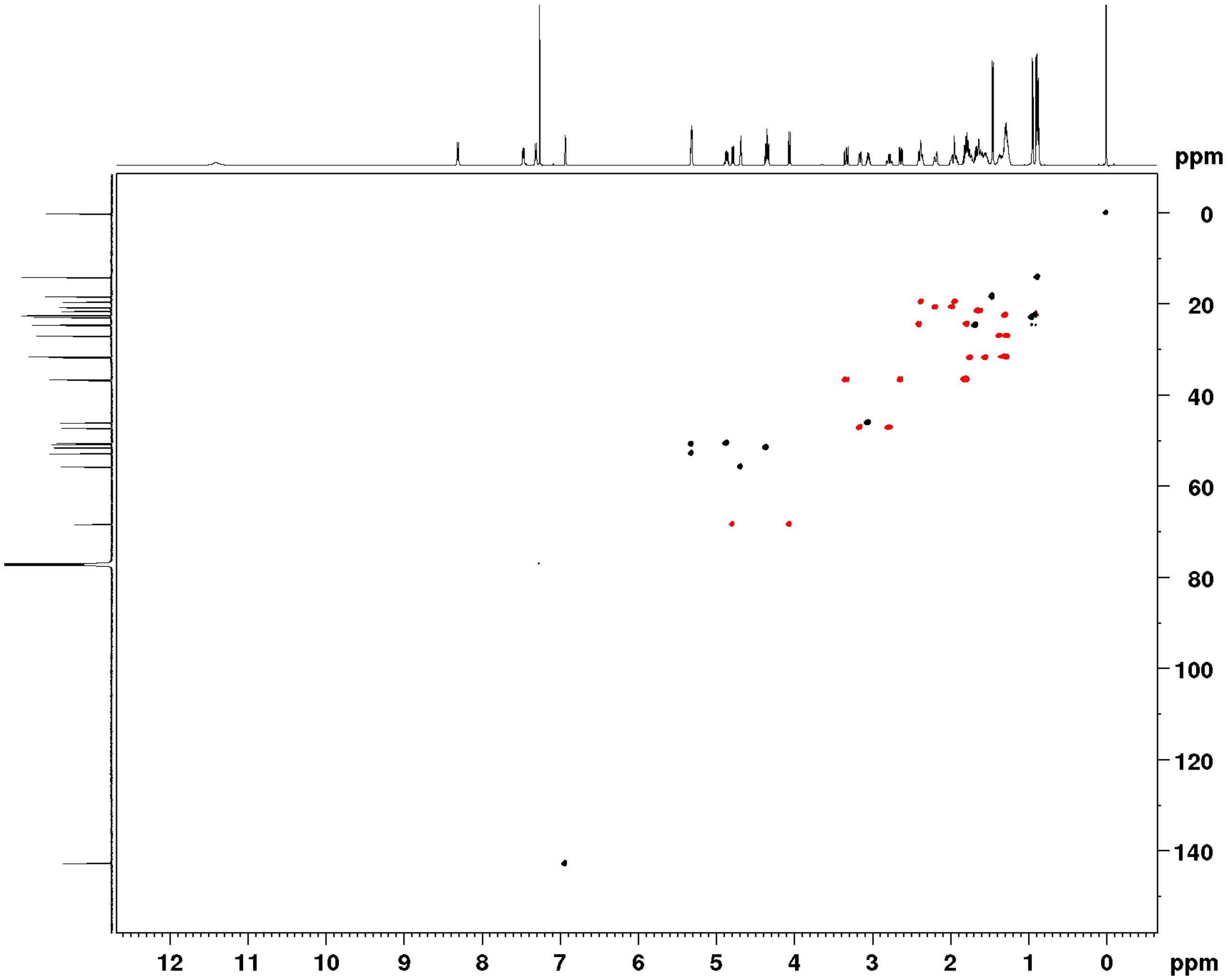


**Figure S14** Lydiamycin A multiplicity-edited HSQC NMR spectrum ($CDCl_3$, 298 K).

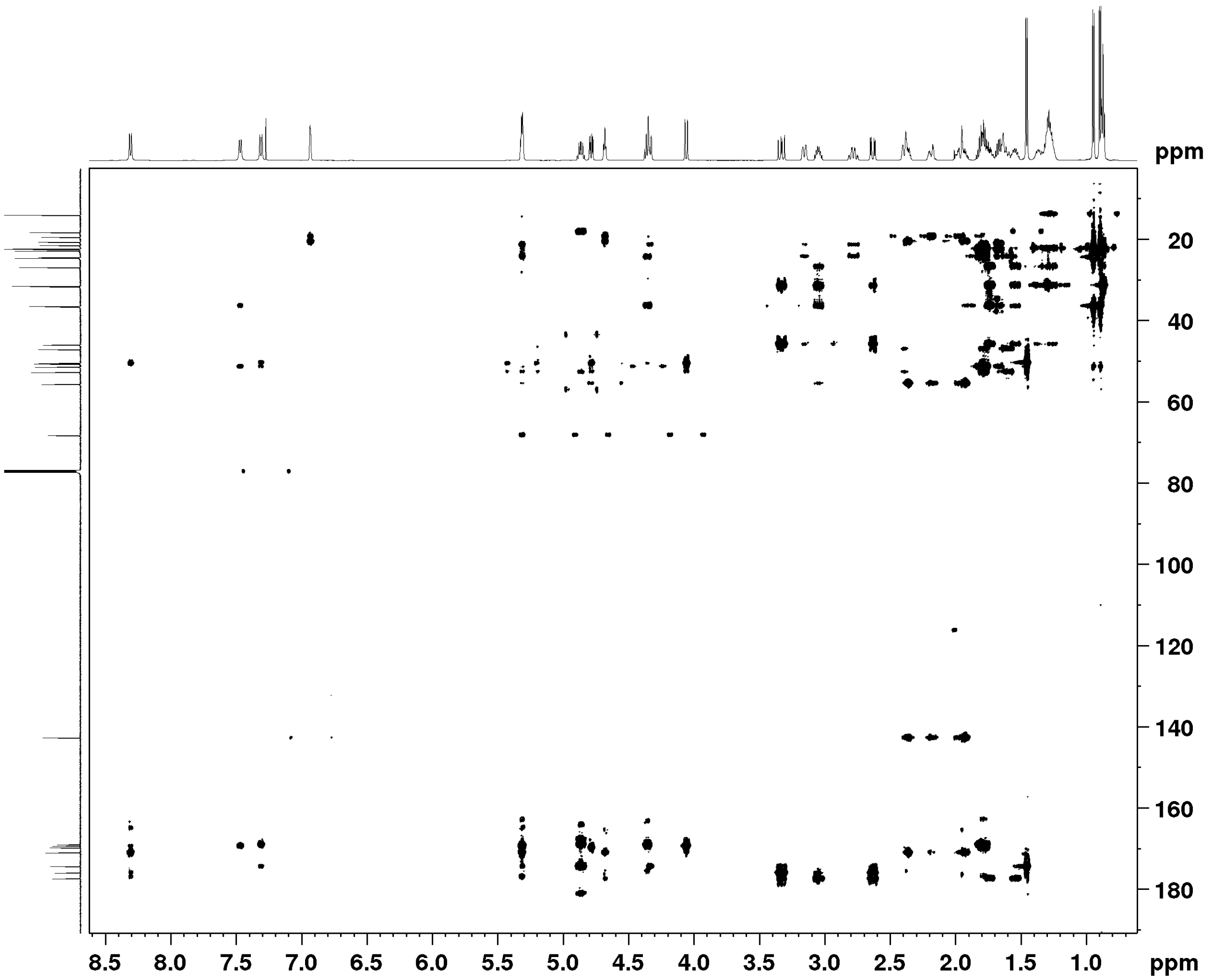


**Figure S15** Lydiamycin A $^{1}H$-$^{13}C$ HMBC NMR spectrum ($CDCl_3$, 298 K).

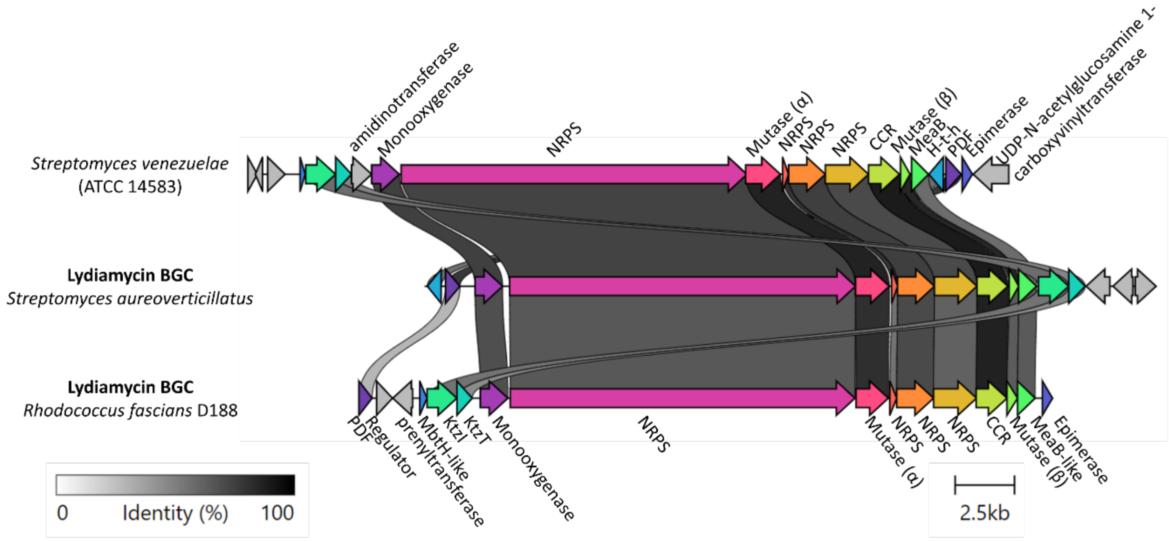


**Figure S16** Comparison of known lydiamycin BGCs with a proposed lydiamycin BGC from *S. venezuelae*. Sequence identity is denoted by greyscale linker. The *S. aureoverticillatus* BGC is described in Libis *et al*.[38]

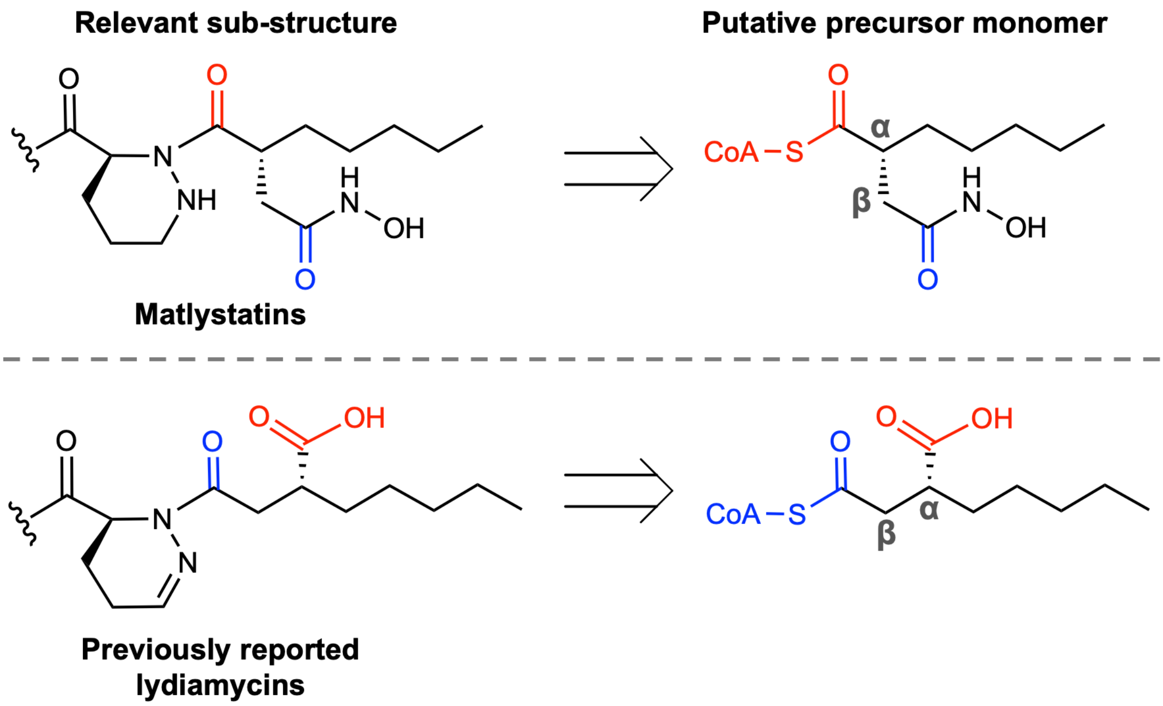


**Figure S17** Schematic of potential structural inconsistency of the published lydiamycin A structure when compared to matlystatin. The alpha-proximal carbonyl groups are highlighted in red and the beta-proximal carbonyl groups are highlighted in blue.

A

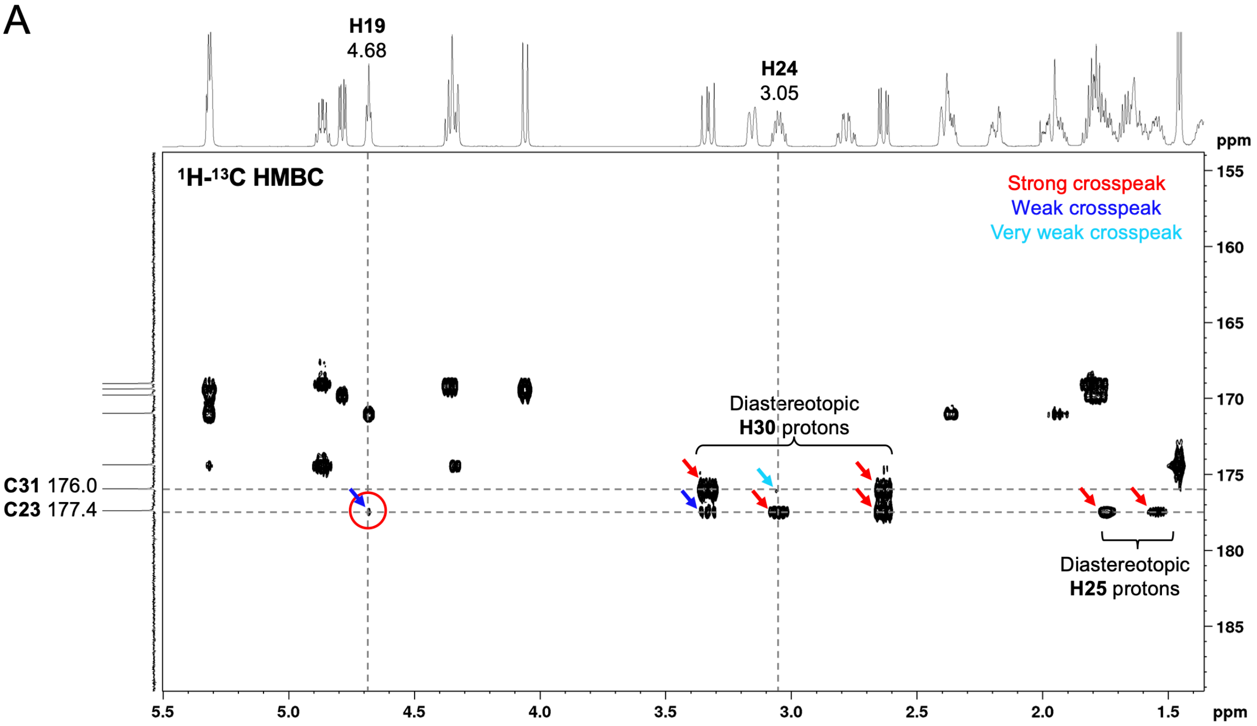


B

| | Observed cross-peaks | | | |
|---|---|---|---|---|
| **Piperazic-Distal Carbonyl C31** ($\delta_C$ 176.0) | | Diastereo $CH_2$ (**H30a**, **H30b**) | CH (**H24**) | |
| **Piperazic-Proximal Carbonyl C23** ($\delta_C$ 177.4) | CH (**H19**) | Diastereo $CH_2$ (**H30a**, **H30b**) | CH (**H24**) | Diastereo $CH_2$ (**H25a**, **H25b**) |

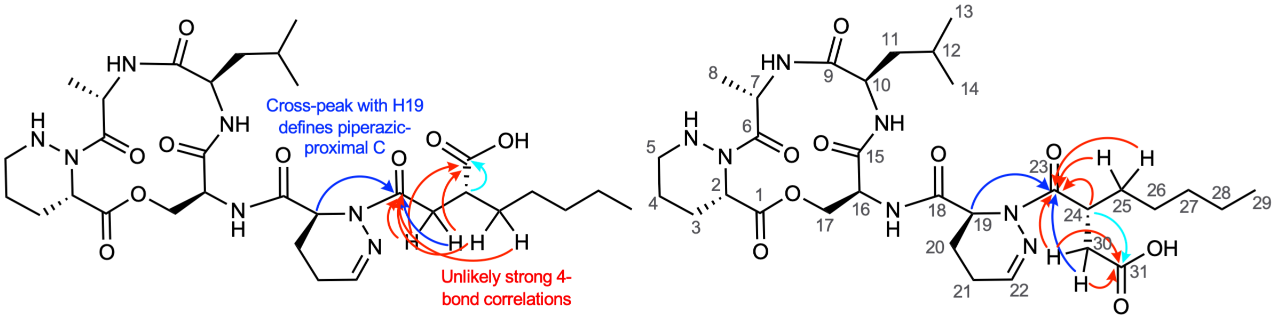


Previously published lydiamycin A structure

Revised lydiamycin A structure

**Figure S18** Analysis of pentylsuccinyl region of HMBC spectrum. A. Selected HMBC correlations for C23 and C31. B. The HMBC cross-peak with H19 enabled the identification of the piperazic-proximal and piperazic-distal carbonyls. The remaining correlations provided unlikely strong correlations given the number of connecting bonds in the proposed published lydiamycin A structure, as well as an absence of correlations between H25 protons and the piperazic-distal carbonyl. A revised structure with an actinonin/matlystatin-like pentylsuccinyl moiety provides correlations that are consistent with the number of bonds between atoms.

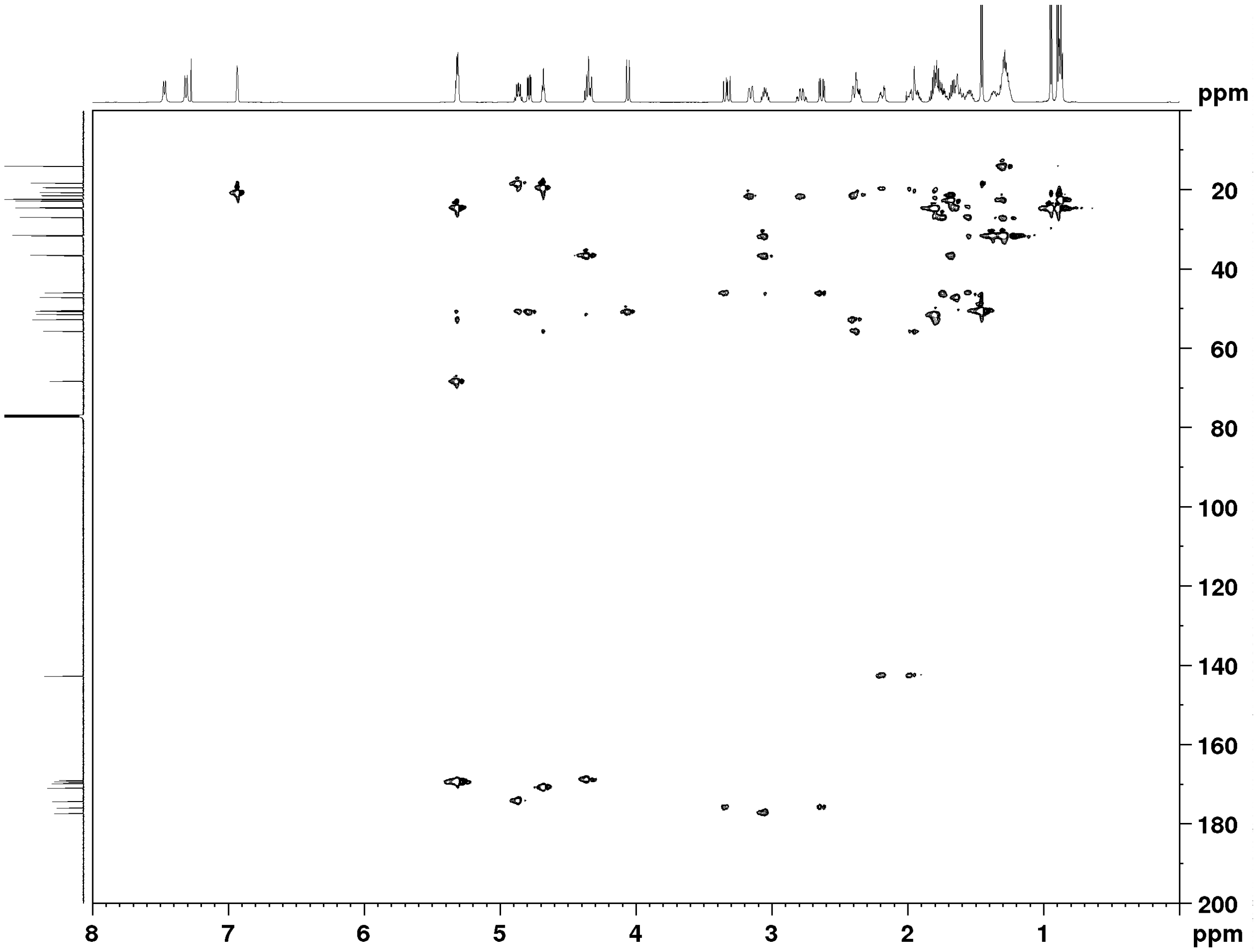


**Figure S19** Lydiamycin A 1,1-ADEQUATE NMR spectrum ($CDCl_3$, 298 K).

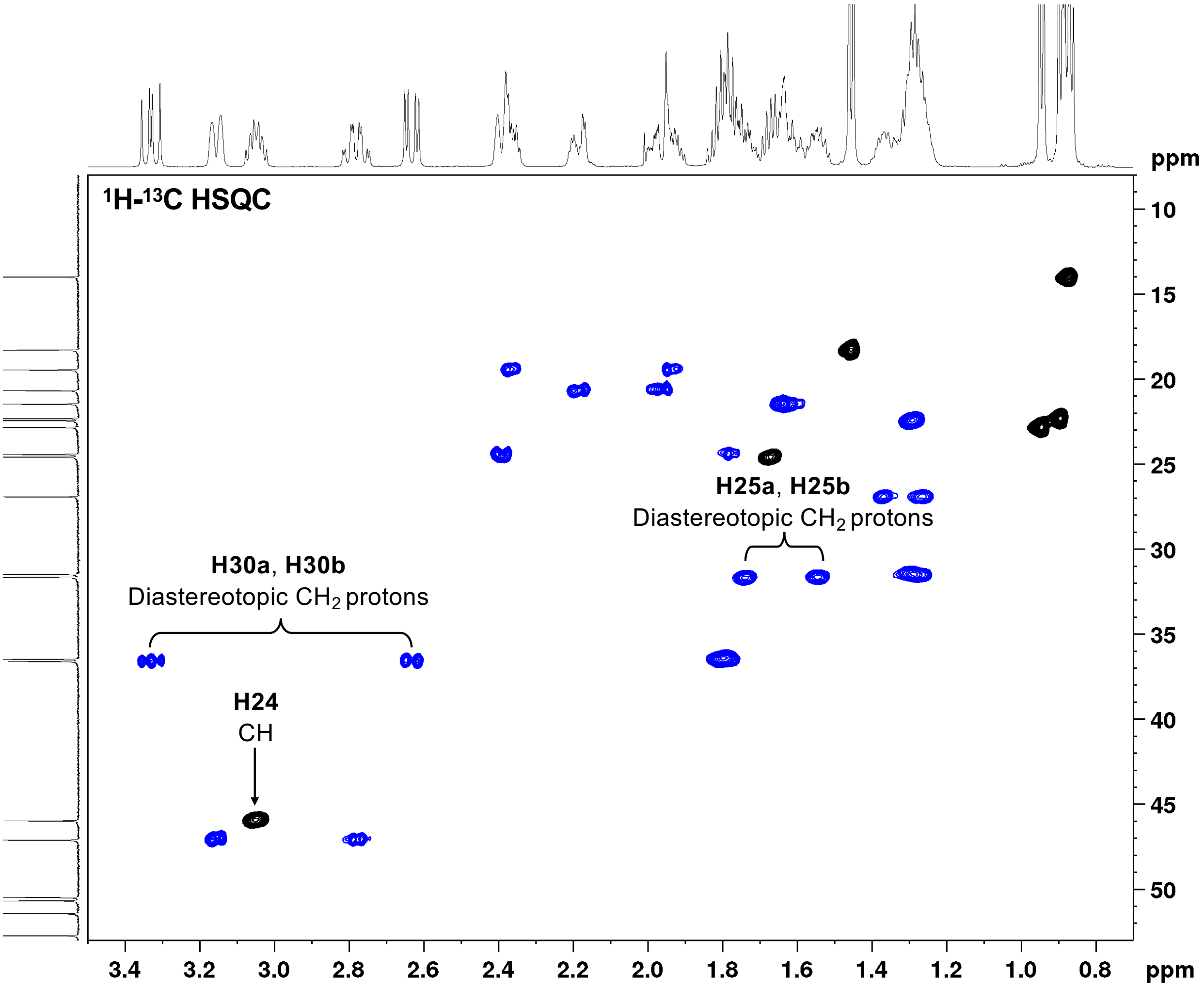


**Figure S20** Region within a multiplicity-edited HSQC NMR spectrum of lydiamycin A. This confirms that the peak at 3.05 ppm (observed in the HMBC data) is from a single proton, which was identified as H24.

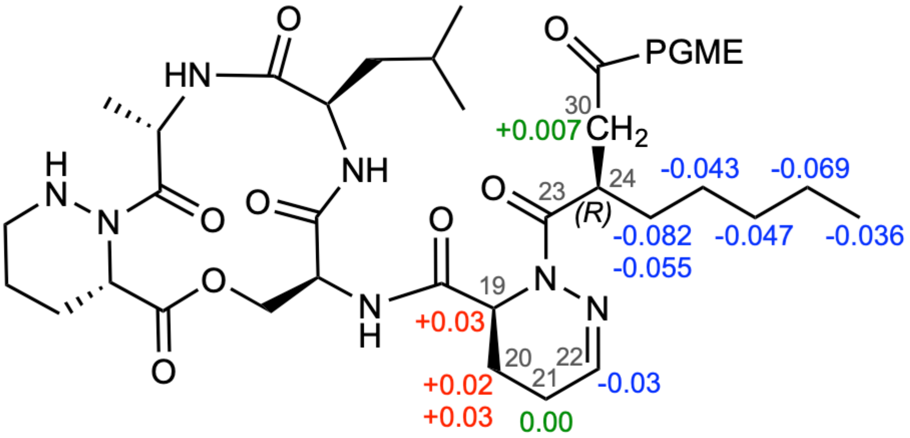


| Proton | Lydiamycin $\delta_H$ | *R*-PGME product $\delta_H$ | *S*-PGME product $\delta_H$ | $\Delta\delta=\delta_{(R)}-\delta_{(S)}$ |
|---|---|---|---|---|
| **19** | 4.95 | 4.96 | 4.93 | +0.03 |
| **20a** | 2.24 | 2.23 | 2.21 | +0.02 |
| **20b** | 1.77 | 1.77 | 1.74 | +0.03 |
| **21a** | 2.15 | Insufficient data | Insufficient data | Insufficient data |
| **21b** | 1.84 | 1.84 | 1.84 | 0.00 |
| **22** | 6.91 | 6.86 | 6.89 | -0.03 |

**Figure S21** Reanalysis of the previously published phenylglycine methyl ester (PGME) derivatisation work published by Hwang and co-workers[36]. The modified Mosher method described by Yabuuchi and Kusumi was applied for using PGME derivatives to determine the absolute configuration of β,β-disubstituted propionic acids[39]. This method involves calculating $\Delta\delta=\delta_{(R)}-\delta_{(S)}$, which meant using the published $\Delta\delta_{S-R}$ (ref. [36]) and inverting the sign of the value. The calculations were also extended to use the published $^1$H NMR for the PGME derivatives to calculate the $\Delta\delta_{R-S}$ values for the dehydropiperazic acid protons 19-22. When the revised lydiamycin A structure is represented with the $CH_2$CO-PGME group with a wedge bond, the protons with $\Delta\delta_{R-S}<0$ are to the right and $\Delta\delta_{R-S}>0$ are to the left of this group. These data are consistent with absolute stereochemistry at C24 of *R*. We note that the PGME plane seems to pass through C21, which has a $\Delta\delta_{R-S}$ of 0.00. Furthermore, we note that the published Δδ value for what we have annotated as C30 is a very small magnitude (+0.007), which is expected as it is also in the PGME plane.

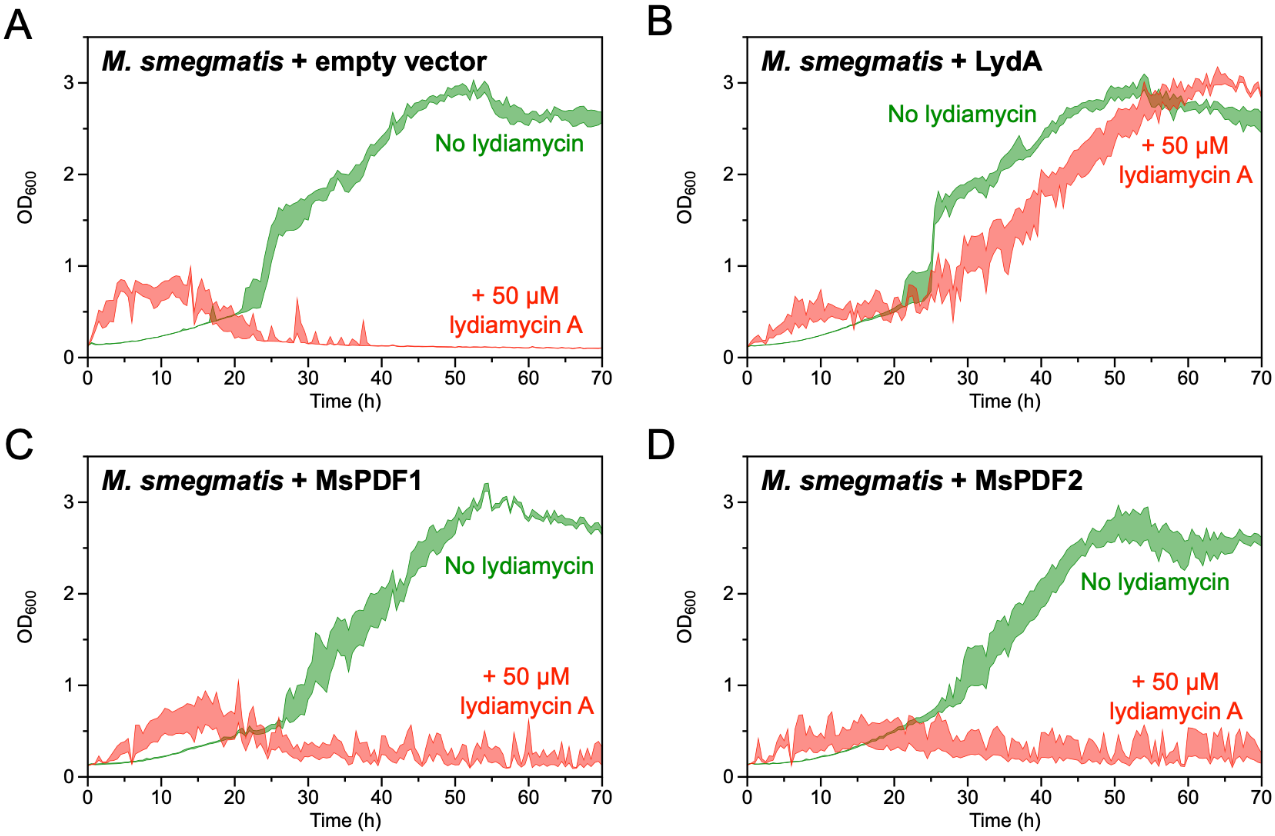


**Figure S22** Assessment of lydiamycin A sensitivity for *M. smegmatis* mc$^2$155 overexpressing its native housekeeping PDF gene (*MsPDF*) compared to LydA. The standard error of the mean is illustrated by the shaded envelope for each strain in the absence or presence 50 μM lydiamycin A (n=6 for each condition). The fluctuations in the curves are a consequence of the non-disperse growth of *M. smegmatis*. A. *M. smegmatis* with empty pJAM2 vector. B. *M. smegmatis* with pJAM2-*lydA*. C and D. Two independent transformants of *M. smegmatis* with pJAM2-*MsPDF*.

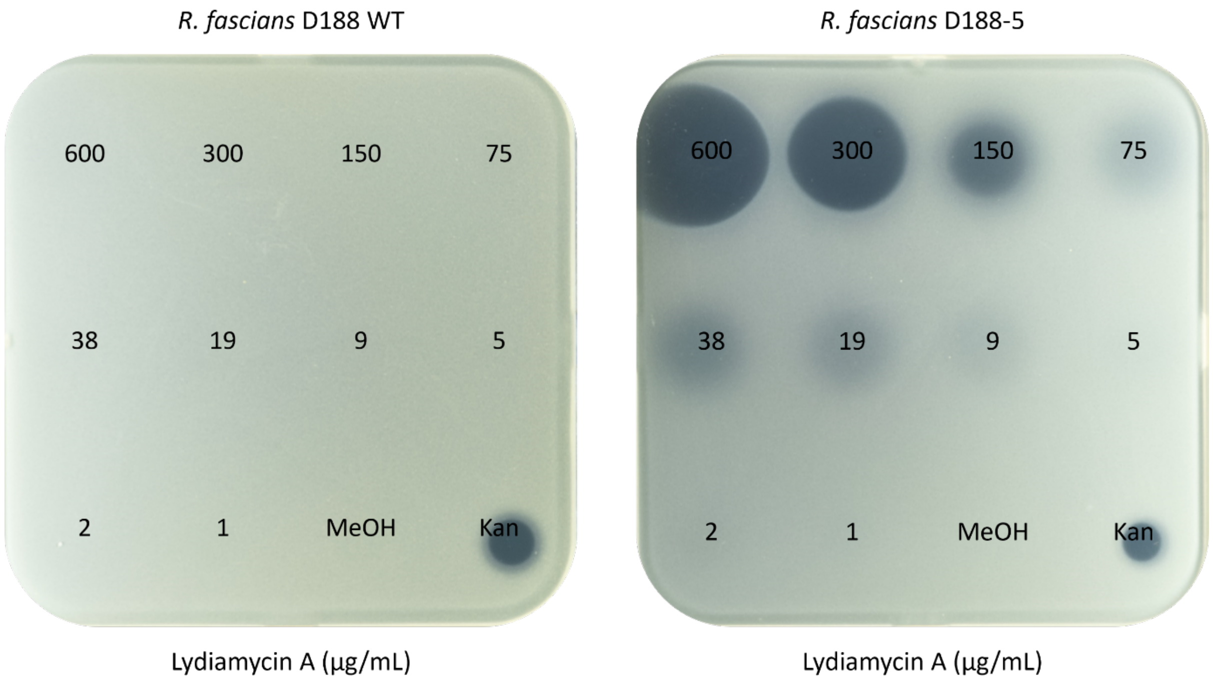


**Figure S23** Spot-on-lawn assays of lydiamycin A (600 to 1 μg/mL) against *R. fascians* WT (left) and plasmid-free *R. fascians* D188-5 (right). Kan = 50 μg/mL kanamycin positive control.

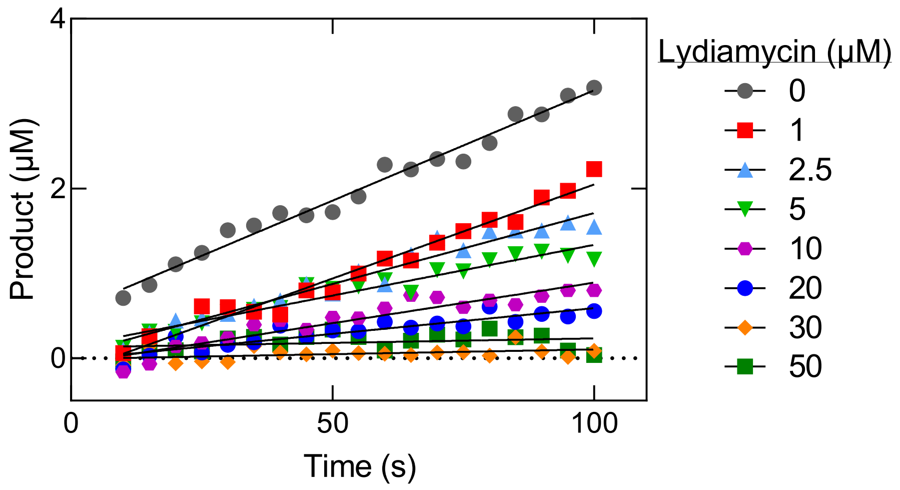


**Figure S24** Examples of the deformylation time courses. fML-pNA (20 μM) was deformylated by *E. coli* PDF (10 nM) in the presence of increasing lydiamycin A concentrations. Lines represent the linear regression used to determine the initial velocity shown in Figure 4.

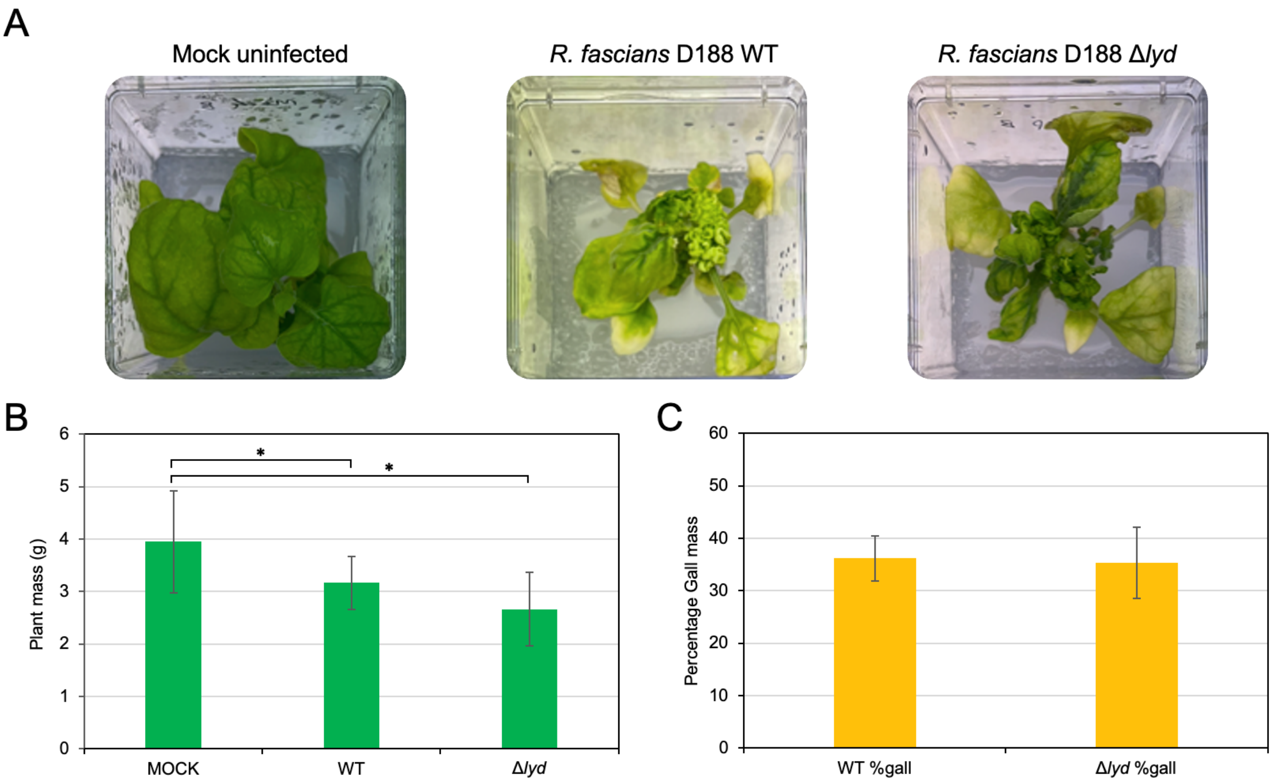


**Figure S25** Infection assays with whole *Nicotiana benthamiana* plants (n = 9 plants each). A. Representative images of infected *N. benthamiana* plants. B. Average fresh plant mass (g +/- standard deviation) of plants 4 weeks post infection with either mock solution (100 mM $MgCl_2$), *R. fascians* D188 WT or *R. fascians* D188 Δ*lyd*. The masses of WT infected and Δ*lyd* infected plants were statistically lower than the masses of uninfected plants (Student's *t*-test: $p$=0.048, 0.024, respectively). There was no significant difference in plant mass between WT infected and Δ*lyd* infected plants. C. Average fresh gall mass in respect to whole plant mass for *R. fascians* D188 WT or *R. fascians* D188 Δ*lyd*.

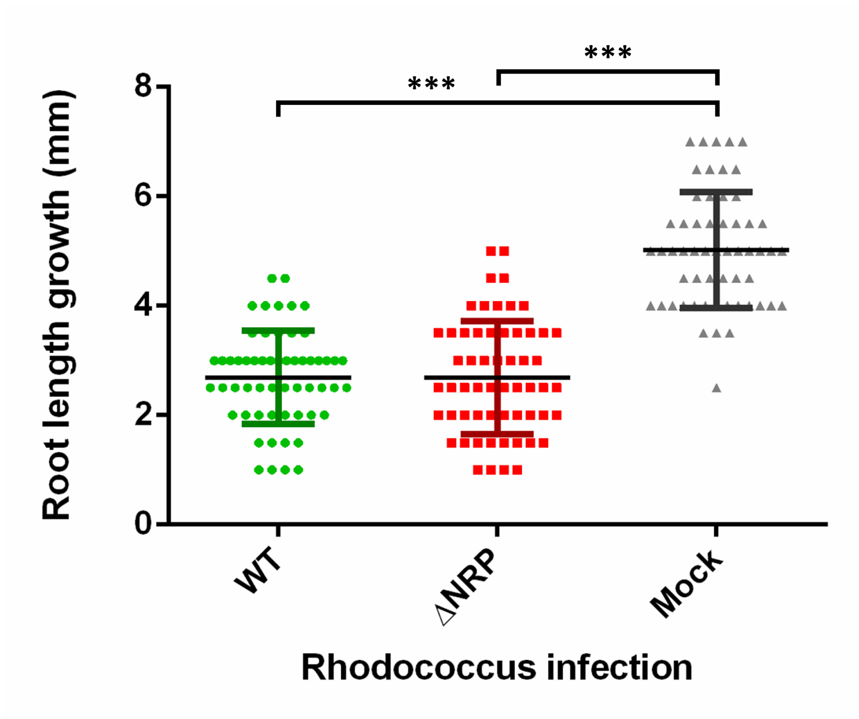


**Figure S26** *N. benthamiana* root length assay with either mock solution (100 mM $MgCl_2$), *R. fascians* D188 WT or *R. fascians* D188 Δ*lyd* (ΔNRP). 60 seedlings were used per condition and root growth was assessed after 7 days. *** = Student's *t*-test significance where p<0.00005.

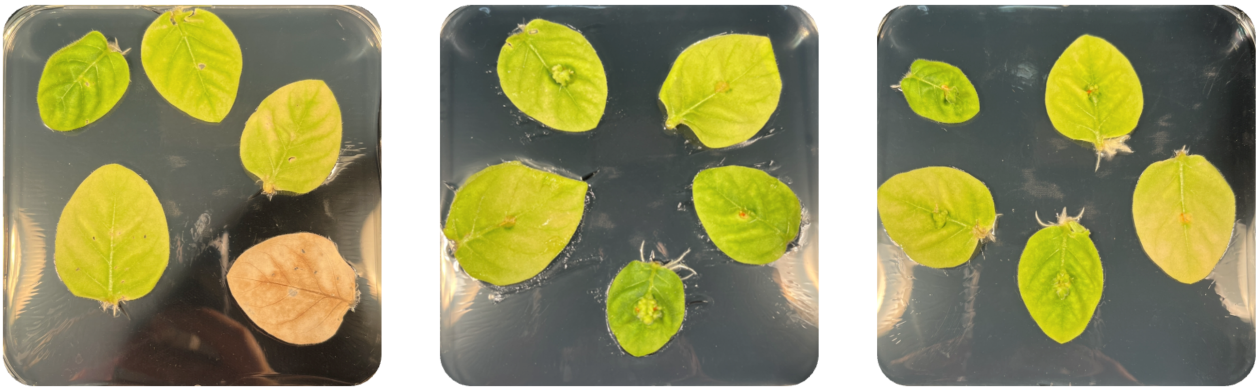

Mock Infection | *R. fascians* D188 WT Infected | *R. fascians* D188 Δ*lyd* Infected

**Figure S27** Representative images from an excised leaf assay with *N. tabacum* infected with mock solution (100 mM $MgCl_2$), *R. fascians* D188 WT or *R. fascians* D188 Δ*lyd* (n = 15 leaves per condition). Leaves were assessed three weeks after infection.

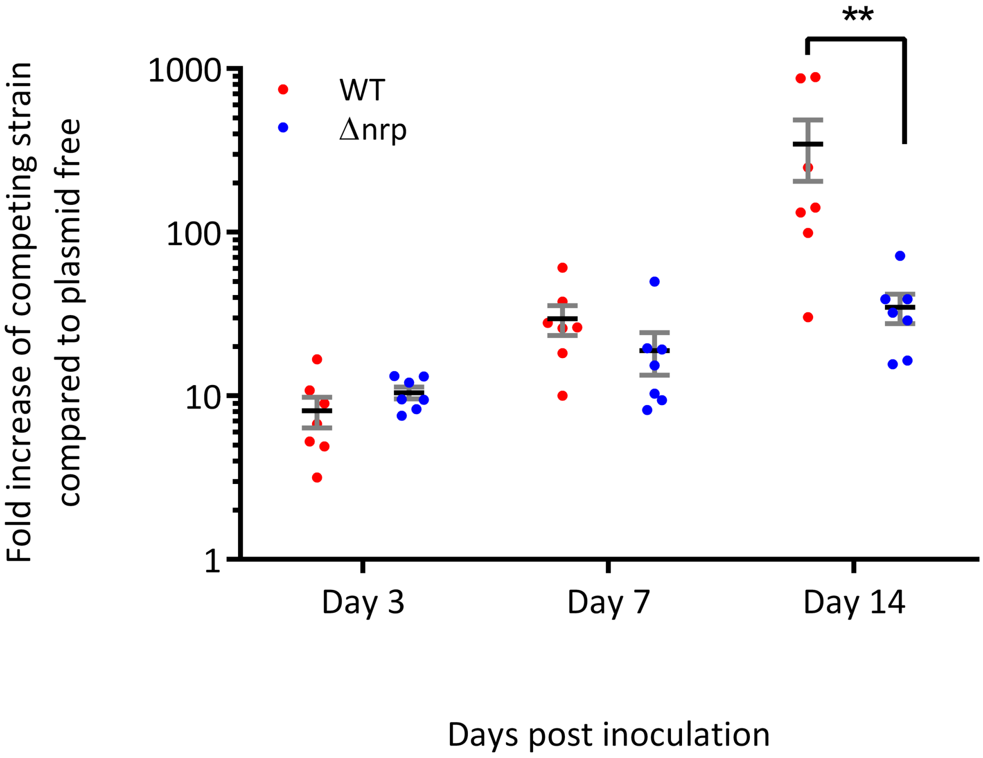


**Figure S28** Competition assay of plasmid-free *R. fascians* D188-5 against WT and Δ*lyd* strains across three time points. Percentage of WT (red) and Δ*lyd* (blue) are plotted as a fold-increase compared to the plasmid-free strain. Individual biological replicates are indicated as dots and the mean of these biological replicates is shown as a black cross-bar. Error bars (grey) indicate the standard error of the mean. A Two-way ANOVA showed a significant interaction between strain and time point ($p<0.05$) with a significant effect of both days post inoculation ($p<0.01$) and competing strain ($p<0.05$). Sidak's multiple comparisons showed a significant difference between WT and Δ*lyd* at day 14 post inoculation ($p<0.01$, **).
